# Let’s make it clear: Systematic exploration of mitochondrial DNA- and RNA-protein complexes by complexome profiling

**DOI:** 10.1101/2023.04.03.534993

**Authors:** Alisa Potter, Alfredo Cabrera-Orefice, Johannes N. Spelbrink

## Abstract

Complexome profiling (CP) is a powerful tool for systematic investigation of protein interactors that has been primarily applied to study the composition and dynamics of mitochondrial protein complexes. Here, we further optimised this method to extend its application to survey mitochondrial DNA- and RNA-interacting protein complexes. We established that high-resolution clear native gel electrophoresis (hrCNE) is a better alternative to preserve DNA- and RNA-protein interactions that are otherwise disrupted when samples are separated by the widely used blue native gel electrophoresis (BNE). In combination with enzymatic digestion of DNA, our CP approach improved the identification of a wide range of protein interactors of the mitochondrial gene expression system without compromising the detection of other multi-protein complexes. The utility of this approach was particularly demonstrated by analysing the complexome changes in human mitochondria with impaired gene expression after transient, chemically-induced mtDNA depletion. Effects of RNase on mitochondrial protein complexes were also evaluated and discussed. Overall, our adaptations significantly improved the identification of mitochondrial DNA- and RNA-protein interactions by CP, thereby unlocking the comprehensive analysis of a near-complete mitochondrial complexome in a single experiment.

## Introduction

Mitochondria are dynamic cell hubs involved in numerous biological processes comprising not only energy conversion but also calcium homeostasis, signal transduction, gene expression, immunity, stress, etc. (1). These organelles contain multiple copies of their own genetic material, i.e. mitochondrial DNA (mtDNA), which encodes a few essential gene products for the oxidative phosphorylation (OXPHOS) pathway, two mitoribosomal RNAs (mt-rRNAs) and a full set of mt-tRNAs (2). To synthesize the mtDNA-encoded proteins, both strands of the circular mtDNA are transcribed into polycistronic mtRNAs. The transcripts are then processed and maturated by multiple mtRNA-processing enzymes and translated by mitoribosomes. Mitochondrial DNA replication and gene expression systems are tightly regulated by specific sets of nuclear-encoded proteins that, upon synthesis by cytosolic ribosomes, are imported into mitochondria. To execute their functions, these proteins interact with mtDNA and mtRNA molecules at specialized membrane-less compartments such as nucleoids (3) and RNA granules (4). While many mtRNA-interacting proteins also, or only, operate outside the confines of RNA granules, a considerable number also have a function typically associated with nucleoids; e.g. during the early steps of mitoribosome biogenesis (5).

Although the molecular roles of the major mitochondrial nucleic acid-interacting proteins have been described, a lot of interactors remain unverified or unknown. Detailed description of such interactions is mandatory for understanding their specific function(s) and integrating them properly into the physiological context. Different strategies are available to explore mtDNA- and mtRNA-protein complexes based on affinity purification, proximity labelling, cross-linking, confocal microscopy, etc. Commonly, these methods require genetic interventions or existence of antibodies/probes/ligands that, on one hand, are time- and material-consuming and, on the other, might influence the stability, quantity and interactions of protein complexes. In addition, these methods do not specifically resolve subpopulations of complexes, a relevant requisite for appropriate depiction of multitasking proteins and their interactions. Incorporation of advanced integral strategies is important to overcome these issues and create accessible options for systematic characterization of mitochondrial nucleic acid-binding protein complexes in different cell types and tissues.

In recent years, a comprehensive mass spectrometry (MS)-based approach known as complexome profiling (CP) has been extensively applied for examination of mitochondrial multi-subunit protein complexes (6). CP is an unbiased method that combines fractionation of native proteins with quantitative MS identification followed by computational data clustering and analysis (7,8). The resulting abundance profiles of proteins that form complexes typically display similar patterns of migration/elution. This method allows the detection of known and potential protein interactors, as well as subcomplexes and assembly intermediates. The standard CP setup includes native gel electrophoresis as protein separation method due to its major advantages, i.e. the low amount of starting biological material needed and better estimation of molecular masses based on its higher resolution.

Despite its benefits, CP has not been applied to systematically investigate nucleoprotein complexes. It has nonetheless been used to study the plant mitoribosomal subunits stabilised by cross-linking (9), leading to identification of new mitoribosome-associated proteins. In human cells, this approach has been restricted for assessing the abundance and integrity of mitoribosomes in mutant and patients cells lines (10–12). CP approaches based on density gradient ultracentrifugation have also been implemented to examine mitoribosomes (13,14). Not surprisingly, CP has not been employed for exploring the nucleoid-associated proteins, because mtDNA molecules coated with tightly-bound proteins are too large to enter standard native gels. Recently, however, analysis of chromatin-enriched fractions separated by blue native electrophoresis (BNE) followed by tandem MS has been performed (15). A large number of well-known chromatin complexes and novel components were identified in this study, which demonstrated that native electrophoresis is suitable for exploring DNA-protein interactions.

As native gel-based CP offers easier experimental handling and the aforesaid advantages, we here present an optimised protocol aimed to investigate protein interactors of the mitochondrial gene expression system. To achieve this, we compared two variants of native electrophoresis and optimised sample preparation conditions, which enhanced the identification of such interactions in human mitochondria. We thoroughly described the composition and features of many mtDNA-/mtRNA-interacting protein complexes detected by this approach, providing a good basis for interpretation of future investigations. Overall, we demonstrate that our CP approach offers a convenient and systematic way of analysing interactions of proteins involved in mtDNA-/mtRNA-related processes. In principle, this method can be used equally well for investigation of other DNA-/RNA-protein interactions regardless their cell location.

## Materials and methods

### Cell culture and treatments

Human embryonic kidney 293 (HEK293) cells (ATCC; CRL-1573) were grown in high-glucose DMEM (Lonza; BE12-604F) supplemented with 10% foetal bovine serum (FBS) (Gibco; 16000044) in a humidified 37 °C incubator at 5% CO_2_. The cell line was routinely tested negative for mycoplasma contamination. To inhibit mtDNA replication and transcription, the medium was supplemented with 30 ng/ml of ethidium bromide (EtBr) for 3 days (72 hours). To further reinitiate mitochondrial DNA and RNA synthesis, the EtBr-containing medium was removed and cells were washed three times with fresh medium. The cells from each plate were transferred into two new plates and incubated for another 24 hours.

### Isolation of mitochondria

HEK293 cells were harvested by resuspension in fresh growth medium, washed with cold phosphate-buffered saline (PBS), pelleted by centrifugation (350 x *g*; 3 min; 4 °C), and resuspended in isotonic mitochondrial isolation buffer (250 mM sucrose, 1 mM EDTA, 20 mM Tris-HCl, pH 7.4, protease inhibitor cocktail SigmaFAST; Sigma; S8820). To isolate mitochondria, the cells were disrupted by applying 15 strokes in a Potter-Elvehjem tissue grinder. The homogenate was diluted with three volumes of isolation buffer and centrifuged (1000 x *g*; 10 min; 4 °C) to remove cell debris and nuclei. The supernatant was transferred to new tubes and centrifuged again to remove the remaining debris. The crude mitochondrial fraction was pelleted from the cleared supernatant (11,000 x *g*; 10 min; 4 °C). Then, the mitochondrial pellet was re-suspended in 0.5 ml of isolation buffer, loaded onto a two-step sucrose gradient (1.5 M/1 M sucrose in 20 mM Tris-HCl pH 7.4, 1 mM EDTA), and centrifuged at 60,000 x *g* for 20 min at 4 °C in a swing-out rotor. The purified mitochondrial fraction was recovered from the interphase between sucrose layers and washed with three volumes of isolation buffer (11,000 x *g*; 5 min; 4 °C). All the above steps were performed on ice using ice-cold buffers. The solutions were prepared using autoclaved Milli-Q water and filter-sterilized, when possible, to minimize ribonuclease contamination. Protein concentration was determined using the Bradford colorimetric assay (Quick Start™ Bradford Protein Assay Kit; Bio-Rad; 5000201).

### mtDNA copy number measurement

The mtDNA levels were estimated as previously described (16). Briefly, total DNA was isolated from cultured cells and analysed by quantitative real-time PCR (qPCR) using primers for cytochrome *b* (MT-CYB, mtDNA) and amyloid precursor protein (APP, nucDNA). The mtDNA content was normalised to the nuclear DNA (nucDNA) content. Fold changes in the relative mtDNA copy number after EtBr treatment and in recovery samples were calculated using the 2^−ΔΔCT^ method.

### Western blotting

Proteins were detected by western blotting as previously described (16). The following antibodies were used for detection: COXII (Abcam, ab110258); PHB1 (Abcam, ab28172); MRPL3 (Abcam, ab39268), FASTKD2 (ProteinTech, 17464-1-AP); MRPS18b (ProteinTech, 16139-1-AP).

### Northern blotting

Northern blot analysis of RNA was performed as previously described (16) with DIG-labelled probes against 12S mt-rRNA, 16S mt-rRNA and 18S rRNA.

### Protein solubilization

Freshly obtained mitochondrial pellets (200 µg protein) were solubilised with water-soluble digitonin (8 g/g protein; SERVA) in ice-cold solubilization buffer (50 mM NaCl, 2 mM 6-aminohexanoic acid, 50 mM imidazole/HCl, pH 7.0, 2.5 mM MgCl_2_, 0.5 mM CaCl_2_) in a total volume of 47 µl. When needed, the solubilisation mixture was supplemented with 150 U of DNase I (Invitrogen; 18047019) or a combination of DNase I and 5 µg of RNase A (Thermo Scientific; EN0531). For the untreated BNE profile, addition of MgCl_2_ and CaCl_2_ was omitted to enable fair comparison with profiles that were obtained following the protocol described in (17). All the solubilised samples were incubated at 10 °C for one hour. The lysates were supplemented with 2.5 mM EDTA to chelate the divalent cations and further cleared by centrifugation (22,000 x *g*; 20 min; 4 °C). The supernatants were recovered and supplemented with either 7 µl of Coomassie blue loading buffer (35% glycerol, 0.15 M 6-aminohexanoic acid, 1.5% Coomassie brilliant blue G-250 (SERVA)) for BNE or 5 µl of Ponceau loading buffer (50% glycerol, 0.1% Ponceau S) for high-resolution clear native electrophoresis (hrCNE).

### Native gel electrophoresis

The samples were separated by either blue or high-resolution clear native electrophoresis using 3-16% polyacrylamide gradient gels according to published protocols (17,18). The preparation of acrylamide mixtures and gel casting procedure were performed as detailed in (17). The electrophoresis running buffers were the following: anode buffer for BNE and hrCNE: 25 mM imidazole/HCl, pH 7.0; cathode buffers for BNE: 0.02% (dark) or 0.002% (light) Coomassie brilliant blue G-250, 50 mM Tricine, 7.5 mM imidazole, pH ∼7.0; cathode buffer for hrCNE: 0.01% dodecyl-β-D-maltoside (DDM), 0.05% sodium deoxycholate (DOC), 50 mM Tricine, 7.5 mm imidazole, pH ∼7.0. After loading the samples, the gels were initially run for 20 min at 100 V and then for ∼4-5 hours at 200 V until the fronts reached the bottom. For BNE, the run was started using the “dark” cathode buffer and later replaced with the “light” one when the front reached down one third of the gel. All native electrophoretic runs were carried out in a cold room (∼6 °C).

### Gel processing and in-gel trypsin digestion

After electrophoresis, the gels were processed as follows: 1-2 h incubation in fixing solution (50% methanol, 10% acetic acid, 100 mM ammonium acetate), 15 min incubation in staining solution (0.025% Coomassie blue, 10% acetic acid), 1 h incubation in 10% acetic acid for destaining, followed by overnight incubation in water. Each gel lane was cut into 56 slices starting from the bottom. Each slice was diced into small pieces and transferred into a 96-well filter microplate (Millipore; MABVN1250) attached to a liquid-collection plate. The filter-plate wells were prefilled with 200 μL of complete destaining solution (50% methanol, 50 mM ammonium bicarbonate (ABC)). The gel pieces were incubated several times with this solution and shaken until the Coomassie blue dye was fully removed.

For in-gel trypsin digestion, the gel pieces were first incubated with 120 µL dithiothreitol (10 mM in 50 mM ABC; 60 min; shaking; RT), followed by alkylation with 120 µL chloroacetamide (30 mM in 50 mM ABC; 45 min; shaking in the dark; RT). The gel pieces were dehydrated in complete destaining solution for 30 min and then dried for 45 min in a Clean Air^®^ horizontal laminar flow cabinet. Then, 20 μL 5 ng/μL sequencing grade trypsin (Promega; V5111) prepared in 50 mM ABC supplemented with 1 mM CaCl_2_ were added to the gel pieces and incubated 30 min at 4 °C. Lastly, 50 μL 50 mM ABC were added to the wells and the gel pieces were incubated overnight at 37 °C in a sealed bag to minimize evaporation. Next, the peptide-containing solutions were collected onto 96-well PCR microplates by centrifugation. The gel pieces were additionally incubated with 50 μL 30% acetonitrile (ACN), 3% formic acid (FA) (30 min; shaking; RT) and the remaining peptides were collected in the same microplates. The peptides were dried in a vacuum concentrator plus (Eppendorf) and resuspended in 20 μL 5% ACN, 0.5% FA. The samples were stored at − 20 °C if not immediately analysed.

All solutions used for in-gel trypsin digestion were prepared with HPLC-grade reagents and solvents. The liquids were removed or collected from the filter-plate by short centrifugation (15-30 s) at 1000 x *g* in a microplate swinging rotor.

### Liquid chromatography and mass spectrometry

The resulting peptides (3 μL) were analysed by tandem mass spectrometry using a Q Exactive mass spectrometer equipped with an Easy nLC1000 system (Thermo Fisher Scientific). The peptides were separated using emitter columns (15 cm length x 100 μm ID x 360 μm OD x 15 μm orifice tip; MS Wil/CoAnn Technologies; ICT015100015-5) filled with ReproSilPur C18-AQ reverse-phase beads of 3 μm particle size and 120 Å pore size (Dr. Maisch GmbH; r13.aq) on 30 min linear gradients of 5-35% ACN/0.1% FA, followed by 35-80% gradient of ACN/0.1% FA (5 min) at a flow rate of 300 nl/min, and a final column wash with 90% ACN (5 min) at 600 nl/min. The mass spectrometer was operated in positive mode switching automatically between MS and data-dependent MS/MS of the 20 most abundant precursor ions. Full-scan MS mode (400-1400 m/z) was set at a resolution of 70,000 m/Δm with an automatic gain control (AGC) target of 1×10^6^_ions and a maximum injection time of 20 ms. Selected ions for MS/MS were analysed with a resolution of 17,500_m/Δm, AGC target 1×10^5^; maximum injection time 50 ms; precursor isolation window 4.0 m/z. The normalised collision energy was set to 30% at a dynamic exclusion window of 30 s. dd-Settings: Minimum AGC of 1×10^3^, intensity threshold 2×10^4^, peptide match preferred and only precursor ions of charges 2, 3 and 4 were selected for collision-induced dissociation. A lock mass ion (m/z = 445.12) was used for internal calibration.

### MS data processing and quantification

The raw MS data files from all the analysed slices for complexome profiling were processed with MaxQuant v.2.0.3 (19). The identified tryptic peptides were matched to the human UniProt reference proteome (ID UP000005640, release date: 28/04/2021). Sequences of known contaminants were added to the search. The default parameters were kept as default with the following changes: enabled “Match between runs” with a 2-minute time window; minimal peptide length = 6, variable modifications = Oxidation (M), Acetyl (Protein N-term), fixed modification = Carbamidomethylation (C); up to two missed cleavages by trypsin were allowed. To quantify the protein abundances for complexome profiling analysis, the intensity-based absolute quantification (iBAQ) values were calculated.

To quantify the changes in total protein abundance between untreated and DNase-treated samples, all raw MS data files from their three replicates were re-analysed with MaxQuant v2.0.3.0. For each profile, the 56 individual data files were set as 56 fractions of the given replicate. The search parameters were the same as aforementioned, except that deamidation (NQ) was included as an extra variable modification and calculation of label-free quantification (LFQ) values was enabled.

### Complexome profiling

The MaxQuant protein groups output file was processed using the postprocessing script for complexome profiling data (GitHub repository: joerivstrien/process_maxquant). In the resulting complexome dataset, the protein groups with score = −2 were filtered out. Two fractions that exhibited aberrant MS runs were omitted from the analysis (CTRL03-50, EtBr_rec-53). To correct for differences between multiple gel runs, the abundance profiles were aligned using the COPAL tool (20). Alignment was performed separately for BNE- and hrCNE-derived profiles (see **Suppl. Data 6)**. The individual profiles were normalised using the sum of iBAQ values of the identified proteins annotated as mitochondrial in MitoCarta 3.0 (21) and/or mitoXplorer 2.0 (22) and/or as “verified/associated” in the Integrated Mitochondrial Protein Index (IMPI) (23), excluding the subunits of OXPHOS complexes I, III, IV, V and mitoribosomes. Calibration curves of apparent molecular masses was performed as described previously (24) using human membrane and water-soluble protein complexes of known masses (e.g., OXPHOS complexes, VDACs, HSP60, OGDH; for details see **Suppl. Data 6**). The normalised and aligned profiles derived from biological replicates of the same condition were averaged; the individual replicates can be found in **Suppl. Data 6**. The resulting abundance profiles were hierarchically clustered with an average linkage Euclidean algorithm using Cluster 3.0 (25). Each subset of profiles was normalised to the maximal iBAQ value across all slices for each protein to generate the relative abundance profiles. For complexes containing a large number of subunits, e.g., OXPHOS complexes or mitoribosomes, we generated single profiles by averaging the iBAQ (protein abundance) values of all their individual components and normalised when desired. The data were visualised as heatmaps and line charts generated in Microsoft Excel.

### Quantification and statistical analysis

The replicates represent samples derived from independent cell cultures that were run on different gels. The fold-changes in protein abundance within individual peaks or profile segments were determined based on the sum of iBAQ values of the selected slices from each replicate. The changes in abundance of multi-protein complexes were determined based on averaged profiles of all detected subunits. An unpaired two-tailed Student’s t-test was used to test the null hypothesis of no changes between the conditions. Statistical significance was determined using a threshold of *p* ≤ 0.05.

For quantification of total protein abundance between untreated and DNase-treated samples, LFQ intensities from each triplicate were normalised using the respective coefficients obtained from COPAL. Data were processed using Perseus 1.6.15.0. Only proteins detected in all three replicates from at least one condition were considered for the analysis. LFQ values were log_2_-transformed and the missing values were replaced by random values drawn from normal distribution (width = 0.3, downshift = 2.5). The log_2_ fold-change (log_2_FC) values were calculated as the difference between log_2_-transformed averaged triplicates. Statistical significance was determined using a threshold of *p* ≤ 0.05 calculated in two-tailed Student’s t-test. For data visualization, negative log_10_-transformed *p* values were plotted against log_2_FC in R-studio using the ggplot2 package.

## Results and Discussion

### Public complexome profiling datasets contain limited data on mitochondrial DNA-/RNA-binding proteins

Nowadays, several human mitochondrial complexome datasets are publicly available, particularly through the ComplexomE profiling DAta Resource (CEDAR) (26). To determine whether the complexome profiling (CP) datasets contain relevant and specific evidence regarding mitochondrial DNA-/RNA-protein interactors, we inspected datasets that used native gel electrophoresis as the protein separation method. In all cases, mitochondrial proteins were solubilised under similar conditions with the mild neutral detergent digitonin prior to separation by blue native gel electrophoresis (BNE). For illustrative purposes, we selected three public representative datasets, namely CRX15 (27), CRX34 (28), and CRX32 (29). The mitochondrial complexome data were obtained from human skin fibroblasts, HEK293 cells, and osteosarcoma 143B cells, respectively.

To test the quality and distribution of protein signals, we first analysed a subset of well-known mitochondrial proteins, i.e. respiratory complexes I (CI), II (CII), III (CIII), and IV (CIV), F_1_F_O_-ATP synthase (CV), mitochondrial porin (VDAC), isocitrate dehydrogenase (IDH) and prohibitin (PHB) (**Figure 1A**). The migration patterns of all these multi-protein complexes were similar among the datasets, so were their apparent molecular masses (*M*_a_), which matched well their theoretical masses (*M*_t_).

**Figure 1.**
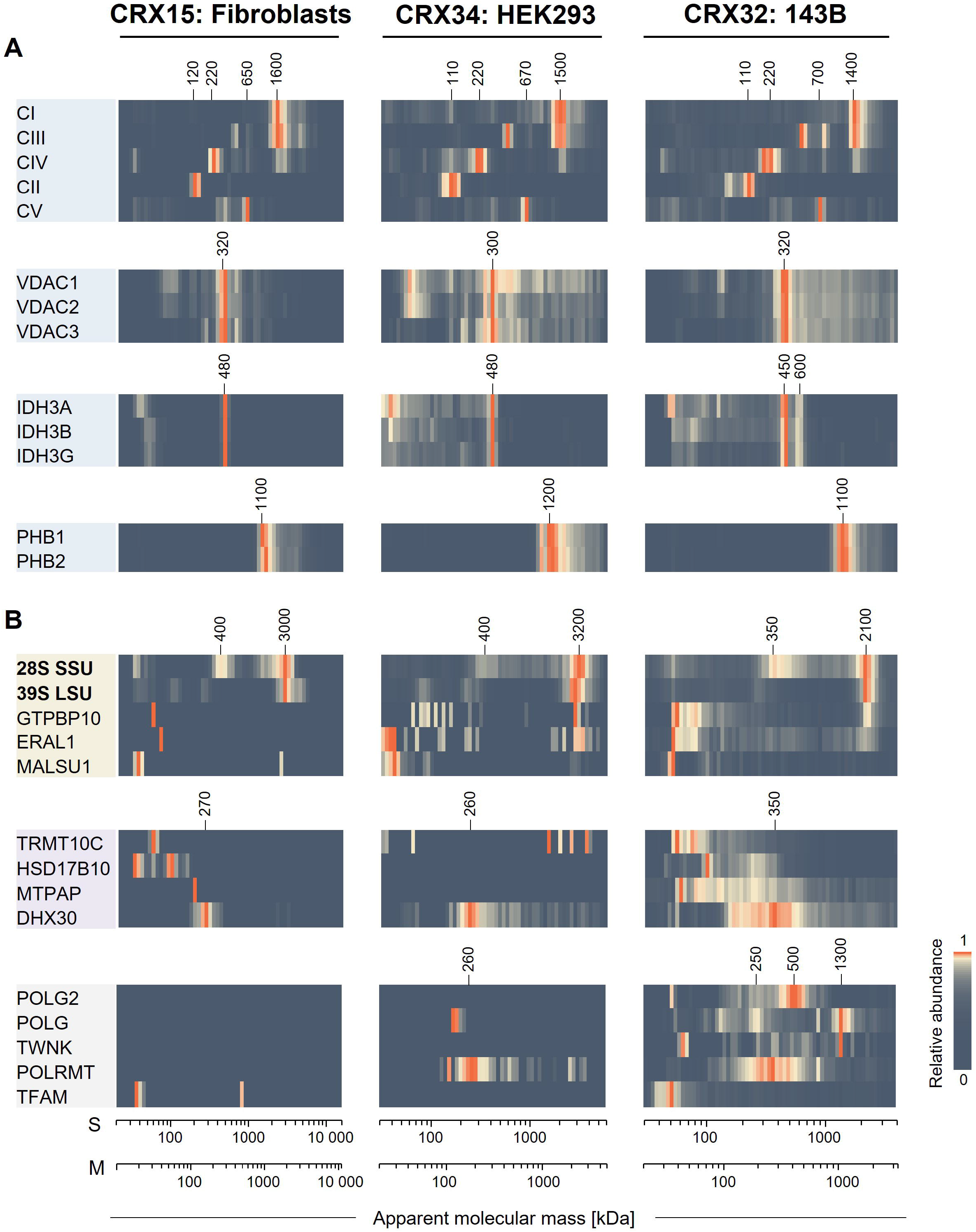
Abundance profiles of selected mitochondrial proteins retrieved from public complexome profiling datasets. Analysis of mitochondrial proteins retrieved from three mitochondrial complexome datasets from human cell lines publicly available in CEDAR. Protein profiles are illustrated as heatmaps showing the relative abundance; i.e. iBAQ values normalised to the maximum value for a given protein in a profile. The migration profiles of the five OXPHOS complexes (CI-CV) and two mitoribosomal subunits (SSU and LSU) were generated by averaging the iBAQ values of all their detected protein components. Specific apparent molecular masses of the complexes are indicated over the heatmaps. **A.** Similar migration profiles of reference protein complexes (blue), i.e. OXPHOS complexes CI-CV, VDAC heterooligomers, IDH complex, prohibitin (PHB), among all three datasets. **B.** Inconsistent detection and migration patterns of mitoribosomes, mitoribosomal assembly factors (blue), mtRNA- (purple) and mtDNA-binding (grey) proteins in the presented datasets. Apparent molecular mass scales are provided for both soluble (S) and membrane (M) proteins.

Next, we looked for mitochondrial proteins known to interact with either mtDNA or mtRNA (**Figure 1B**). Regardless of their abundance, proteins involved in mitochondrial replication or gene expression were inconsistently identified in these datasets. We selected representative proteins that were detected in at least one dataset (**Figure 1B**). In contrast to the reliable detection shown in Figure 1A, abundance and migration patterns of mtDNA-/RNA-binding proteins varied considerably. The protein components of the 28S small (SSU) and 39S large (LSU) mitoribosomal subunits showed similar migration patterns in all cases. Yet, their *M*_a_ differed by ∼1 MDa between the fibroblasts or HEK293 and 143B profiles. Additionally, some of the SSU components accumulated in a sub-assembly at ∼400 kDa. Whereas the mitoribosome-interacting proteins ERAL1 and GTPBP10 were only detected at low *M*_a_ in fibroblasts dataset, small amounts of both proteins comigrated with SSU and LSU in datasets for HEK293 and 143B. The LSU-assembly factor MALSU1 was mainly found at low *M*_a_ with an exception of a portion comigrating with mitoribosomal subunits in the dataset for HEK293. Mitochondrial replication, transcription and mtRNA-processing factors (TFAM, Twinkle, POLG, POLG2, POLRMT, MTPAP, RNase P subunits) were properly identified only in 143B dataset where they displayed miscellaneous migration patterns nonetheless.

Based on this analysis, the identification of abundant and stable multi-protein complexes using BNE-based CP is mostly consistent and comparable. Contrarily, identification of known mtDNA-/RNA-interacting proteins is less reproducible and, in some cases, arguable.

### Detection of mtDNA-/RNA-interacting proteins was improved by using high-resolution clear native electrophoresis in the complexome profiling setup

Separation of macromolecular complexes by native gel electrophoresis and therefore their correct detection by CP can be affected by many factors, e.g., amount of detergent, ionic strength, percentage of polyacrylamide gradient, etc. In fact, some protein-protein interactions are better preserved by using high-resolution clear native electrophoresis (hrCNE) (30,31), i.e. a milder method than BNE, in CP setups. Here, we sought to test whether hrCNE might preserve the stability and facilitate the detection of mtDNA-/mtRNA-protein complexes. Originally, the hrCNE protocol was developed as a “dye-free” alternative to BNE for in-gel fluorescence and enzyme detection assays (18). The Coomassie blue dye is neither added to the samples nor the running buffers. Instead, to impose a negative charge shift on proteins, enable separation by size, keep membrane proteins solubilised and improve resolution of the bands, the cathode running buffer is supplemented with a low concentration of DOC (anionic detergent) and DDM (neutral detergent).

To this aim, we generated complexome profiles of mitochondria isolated from HEK239 cells using both BNE and hrCNE protocols. To ensure the best sample integrity, we performed mitochondria isolation and protein solubilization using ribonuclease (RNase)-free materials and avoided freeze-thawing of the samples.

In the resulting dataset, we observed substantial differences between the complexome profiles obtained with BNE and hrCNE. In the hrCNE-based profile, mitochondrial nucleic acid-interacting proteins had more defined profiles and were coherently clustered in contrast to the BNE-based profile **(Suppl. Data 1)**. To illustrate this, we compared the migration profiles of several well-known mitochondrial translation, mtRNA-processing and transcription factors, as well as mtDNA-binding proteins (**Figure 2A**). Mitoribosome assembly factors (e.g., MTG1, MTG2, MALSU1, NOA1) and rRNA-modifying enzymes (e.g., RPUSD3, MRM1, MRM2) appeared to migrate at higher *M*_a_ in hrCNE gels, and displayed better comigration with mitoribosomes (**Figure 2B**). Various RNA-binding proteins (RBPs), such as mt-mRNA maturation enzymes, mtRNA helicases and transcription factors, which displayed smearing patterns in the BNE gel, were found to peak at high *M*_a_. The mtDNA-binding proteins TFAM, POLG2, and Twinkle also migrated at a higher *M*_a_ in hrCNE gels, although they were detected at relatively low abundance.

**Figure 2.**
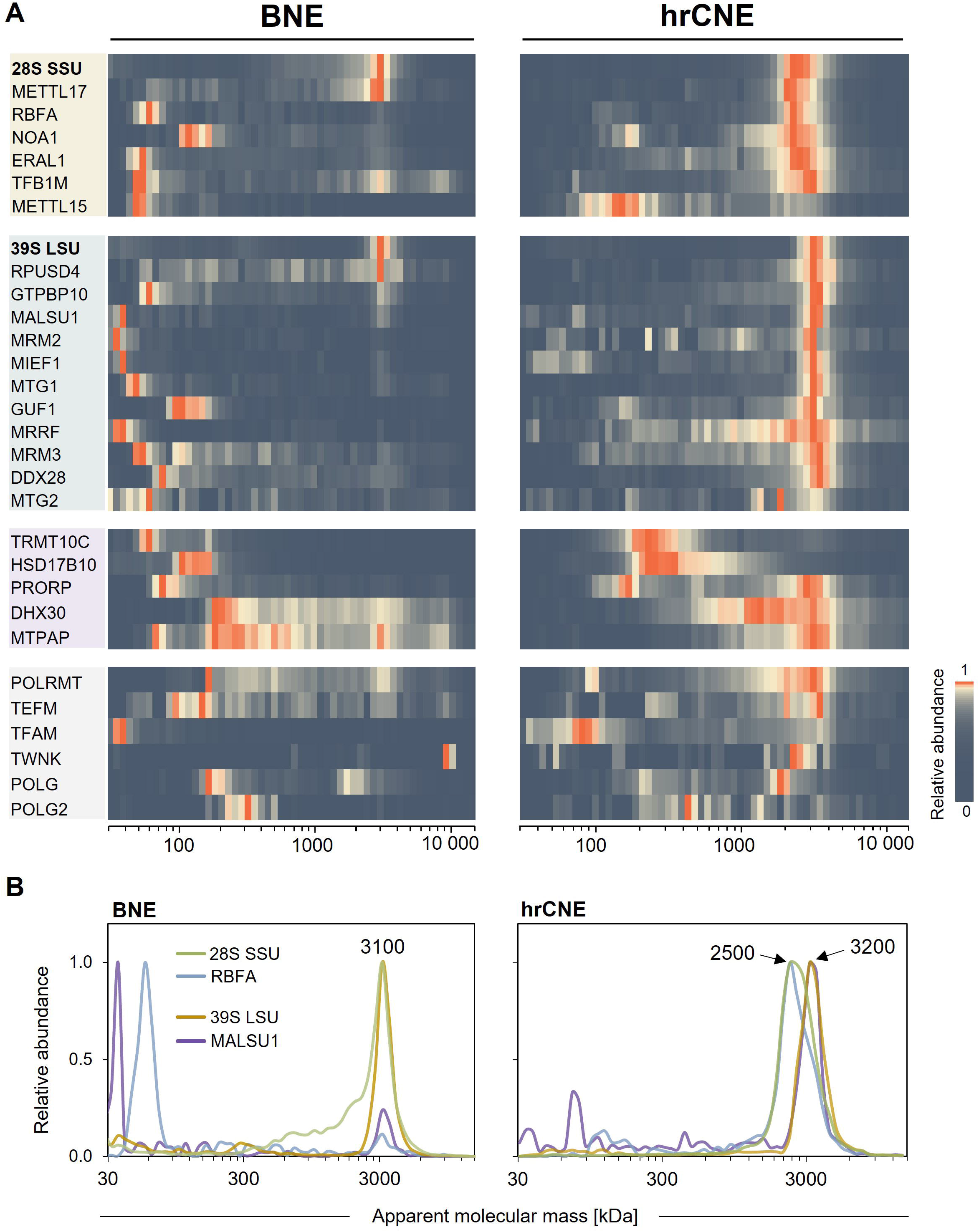
Comparison of mtDNA-/RNA-interacting proteins identified in BNE- and hrCNE-based complexome profiles. A. Mitoribosome-interacting proteins (yellow – SSU, green – LSU) and various mtRNA-processing enzymes (pink), transcription and replication factors (grey) migrate at higher *M*_a_ in hrCNE gels when compared to BNE gels. Heatmaps show relative abundances that represent iBAQ values normalised to the maximum value for a given protein separately for the BNE and hrCNE profiles. **B.** An example of the better comigration of mitoribosome assembly factors (RBFA – SSU, MALSU1 – LSU) with the respective subunits in hrCNE compared to BNE. The relative abundance profiles are illustrated as line charts. The profiles of SSU and LSU correspond to the average of iBAQ values of all their respective detected MRPs. Apparent molecular masses of the main populations are shown next to the peaks in kDa.

In addition, we noticed that the integrity of the mitoribosomal subunits was better in our BNE profile compared to, for example, the published complexome dataset of HEK293 mitochondria (CRX34) (**Figure 3**). Here, the MRPs that migrated at low *M*_a_ were the ones directly interacting with mt-rRNAs on the periphery of the mitoribosome; thus, largely dependent on proper RNA stability. These interactions were likely better preserved in the BNE profile for which we used freshly isolated samples and RNase-free buffers. Notably, however, the mitoribosomal subunits displayed their best integrity when separated by hrCNE, implying that this variant is more suitable to study these ribonucleoproteins. In the hrCNE-based profile, LSU and SSU migrated in two separate peaks in contrast to BNE, reflecting the better resolution of the dye-free variant and/or preservation of additional associated components.

**Figure 3.**
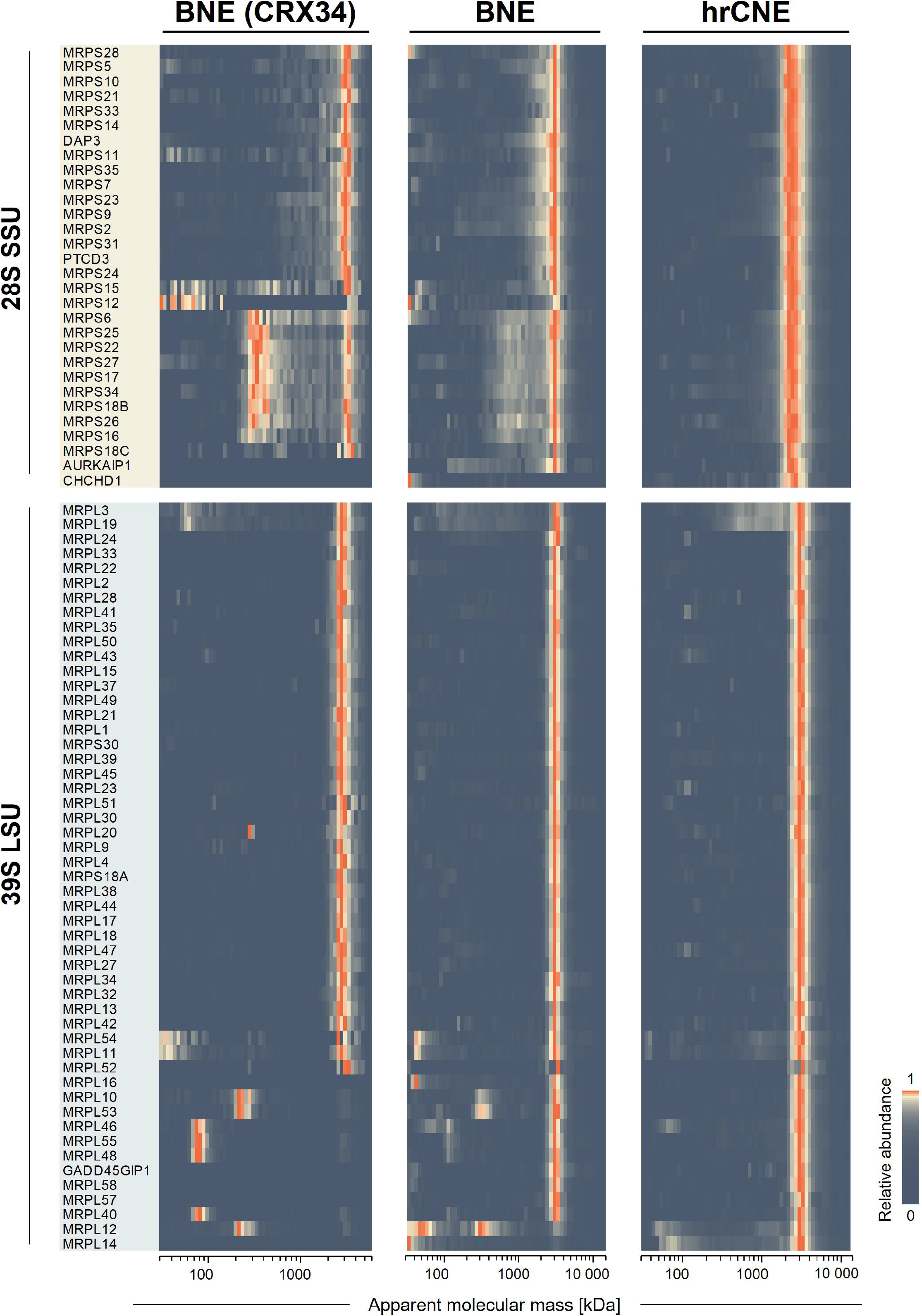
Comparison of mitoribosomal proteins (MRPs) identified in BNE- and hrCNE-based complexome profiles. Heatmaps show the relative abundances representing iBAQ values normalised to the maximum value for a given protein separately for each profile. In the selected BNE-based complexome dataset (CEDAR: CRX34), the mitoribosomal subunits are observed as partially disintegrated. This effect is less pronounced when samples were separated by BNE using RNase-free buffers and freshly isolated mitochondria. Note that mitoribosomal subunits are best preserved when samples are separated by hrCNE using RNase-free buffers and freshly isolated mitochondria.

To test whether the quality of the hrCNE-based CP was overall similar to that of BNE-based CP, we compared the profiles of a subset of reference protein complexes **(Suppl. Figure S1**). The migration profiles of mitochondrial OXPHOS complexes, mitochondrial porin (VDAC), prohibitin, and MICOS-MIB complex were very similar in both datasets and their peaks matched well their reported molecular masses. The mtRNA-binding complex composed of LRPPRC and SLIRP proteins (32) was also properly detected by both approaches. According to what was reported before (29), this complex was found migrating at 200-250 kDa (Suppl. Figure S1). However, detection of some protein complexes was improved by using hrCNE. For example, the ∼4.3 MDa oligomeric 2-oxoglutarate dehydrogenase (OGDH) complex was better preserved in hrCNE gels, as a higher proportion of subunit DLD comigrated with its other three subunits (OGDH, DLST and MRPS36) (33). The membrane proteins ATAD3A and ATAD3B involved in mitochondrial cristae organisation (34,35) comigrated at ∼2 MDa in hrCNE gels instead of ∼1 MDa in BNE gels, indicating the presence of additional interactors preserved in the first case. On the other hand, the isocitrate dehydrogenase (IDH) displayed less defined peaks after separation by hrCNE, reflecting a particular sensitivity of this complex to DOC and/or DDM. Overall, we did not find any notable loss of data quality when using hrCNE for CP.

Taken together, these results demonstrate that the identification of mitochondrial nucleic acid-interacting protein complexes, especially RBPs, is robustly improved after separation by hrCNE without compromising the detection of other protein complexes. Such drastic improvement is likely due to omission of the negatively charged Coomassie blue dye during electrophoresis, which may affect interactions relying on electrostatic forces. The latter are particularly common in contact sites of protein-nucleic acid interactions (36).

### Digestion of DNA improves detection of mtDNA-binding protein complexes by hrCNE-based complexome profiling

After the considerable improvement in the detection of mtRNA-interacting proteins by hrCNE-based CP, we sought to achieve better and reproducible identification of mtDNA-binding proteins. Clearly, the tightly associated proteins are not able to enter regular native gels, as the size of the mtDNA alone (double-stranded circular ∼16.6 kb molecule) is ∼10 MDa. To overcome this limitation, we applied enzymatic digestion of DNA to allow dissociation of such proteins and their entrance in the gel matrix. A similar strategy has been successfully implemented to study chromatin-interacting protein complexes (15). In our experiment, mitochondria were incubated with bovine deoxyribonuclease I (DNase I) during the digitonin-solubilization step and further separated the proteins by BNE and hrCNE in parallel with untreated samples. For simplicity, we will refer to DNase I as DNase hereafter.

In the resulting BNE- and hrCNE-based complexome profiles **(Suppl. Data 2)**, the intensities of several well-characterised mtDNA-binding proteins increased after DNase treatment. However, in hrCNE, additional subpopulations were observed for some of these proteins (**Figure 4**). The total intensity of TFAM (*M*_t_ ∼29 kDa), the mitochondrial transcription and mtDNA-packaging factor (37), was ∼5.5-fold higher in the treated samples in both protocols. While in BNE this protein migrated as a single peak at ∼30 kDa, it distributed in two major peaks at ∼80 kDa (likely representing TFAM oligomers or TFAM with fragments of DNA, as its other known interactors were not found in that range) and at ∼3.2 MDa in hrCNE. Based on the large size and the smeared pattern, the latter may represent a complex with partially digested mtDNA. In a similar case, the subunits of the DNA polymerase gamma (Polγ) (38), composed by a catalytic subunit, POLG (*M*_t_ 140 kDa), and an accessory subunit, POLG2 (*M*_t_ 55 kDa), peaked at ∼500 kDa. This fraction appeared only after DNase treatment in hrCNE, but not in BNE (**Figure 4**). This peak fits well the size of the Polγ holoenzyme complex structure resolved by cryo-electron microscopy consisting of two POLG and four POLG2 copies (∼460 kDa) (39). The other observed peaks likely represent mono- and oligomeric forms of the subunits, possibly including other interactors (see below). Altogether, these results imply that, first of all, digestion of DNA prior to protein separation by native electrophoresis expectedly can enhance the detection of mtDNA-interacting proteins by CP, and, secondly, that hrCNE is again a better option for preserving the integrity of these macromolecular complexes.

**Figure 4.**
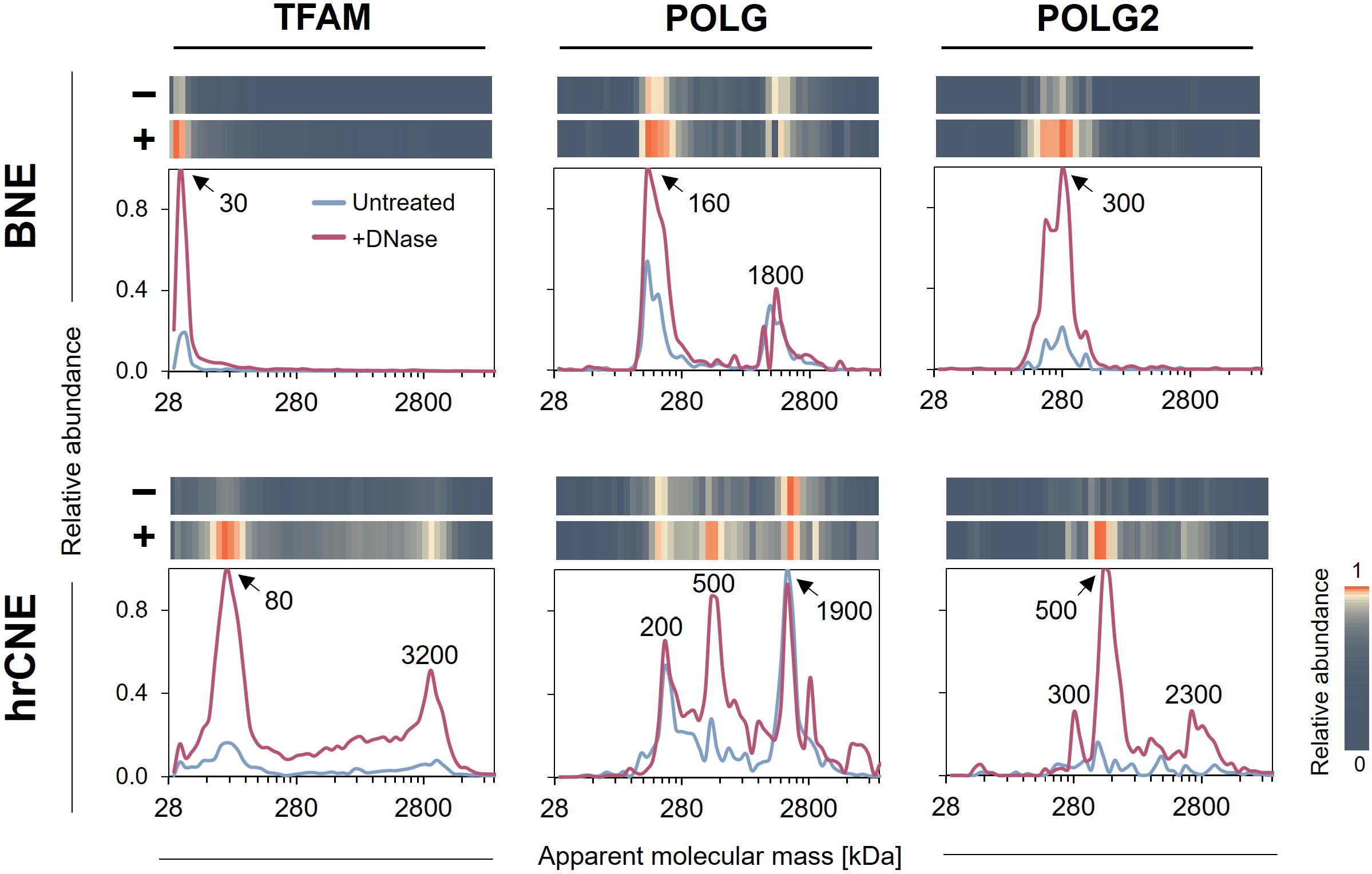
Comparison of mtDNA-binding proteins detected in untreated and DNase-treated samples by BNE- and hrCNE-based complexome profiling. Heatmaps and line charts show migration patterns of TFAM, POLG and POLG2 in the untreated control (-) and +DNase (+) profiles using BNE and hrCNE as protein separation methods for CP. The relative abundances represent iBAQ values normalised to the maximum value for a given protein in each pair (Untreated/+DNase) of profiles separately for BNE and hrCNE. Apparent molecular masses of the main populations are shown over the peaks in kDa.

As it proved to be more reliable than BNE, we performed hrCNE-based CP analysis in triplicate to evaluate the consistency of the DNase treatment **(Suppl. Data 2)**. We observed an increased abundance and/or extra peaks for a number of mtDNA transcription, replication and maintenance factors (40) after DNase treatment (**Fig. 5A**). An especially pronounced effect was detected for TFAM, Polγ, mtSSB, Top3α, and, to a lesser extent, for TEFM and MGME1.

**Figure 5.**
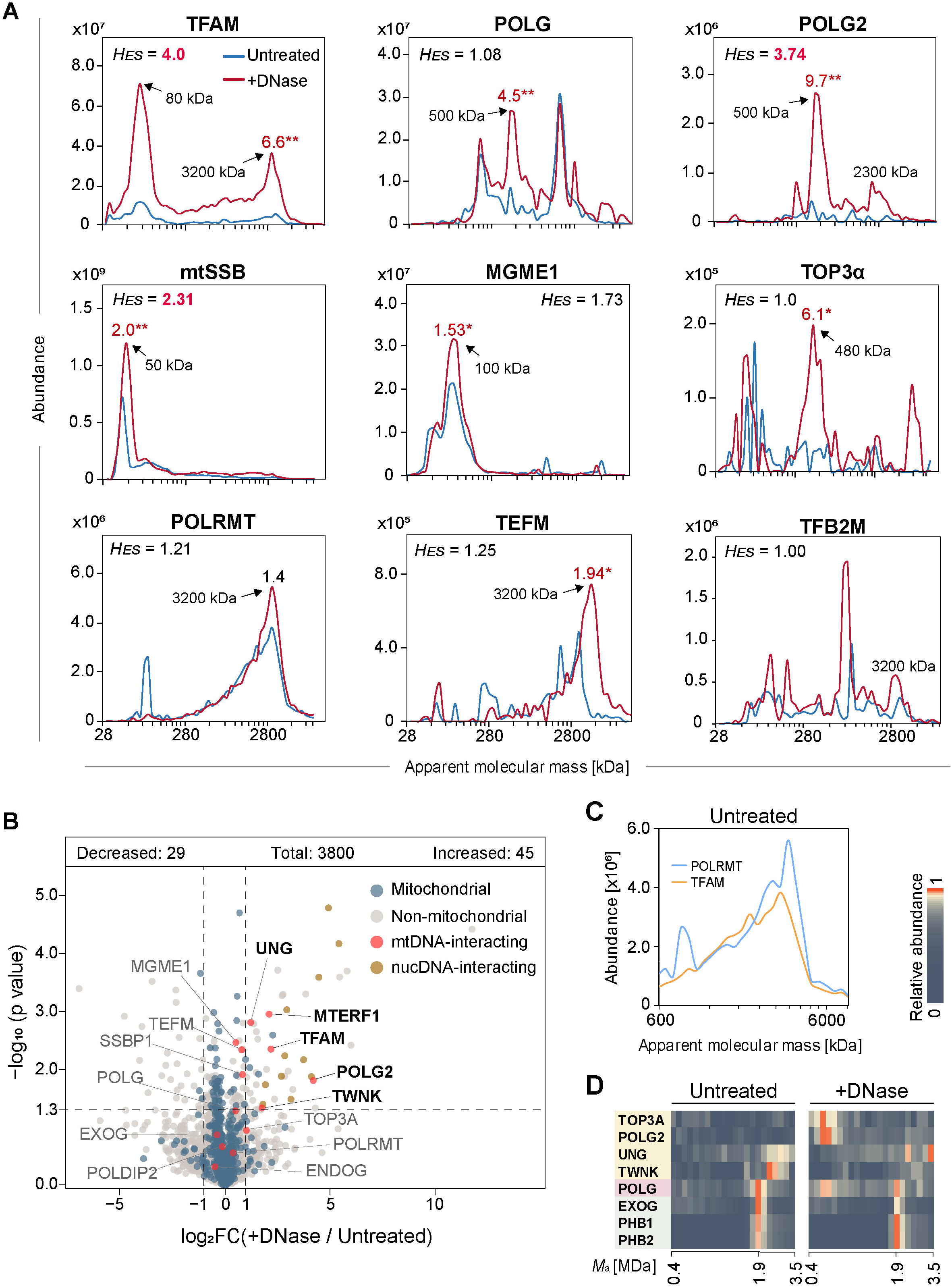
Improved detection of mtDNA-binding proteins after DNA digestion and using hrCNE-based complexome profiling. A. Comparison of abundance profiles of selected mtDNA-binding proteins in untreated and +DNase conditions. The apparent molecular masses of the populations that appeared after treatment are shown over the peaks or with arrows. Hausdorff effect sizes (*H_ES_*) values are shown above the charts, *H_ES_* > 2 values are highlighted in red and indicate substantial changes of the migration profiles. The intensity fold changes within individual peaks, calculated based on sum of iBAQ of 3-5 slices around the peak, are shown above the corresponding peaks. Statistical significance was defined using a two-tailed Student’s t-test (n = 3, \**p* < 0.05, \*\**p* < 0.01). Each profile represents the average of three independent profiles. **B.** Volcano plot depicting negative log_10_-transformed *p* values (two-tailed Student’s t-test; n = 3) plotted against log_2_-transformed fold changes of averaged protein abundances (LFQ intensities). Proteins with −1 ≥ log_2_FC ≥ 1 and *p* ≤ 0.05 were considered affected by the DNase treatment. The thresholds are shown with dashed lines. Proteins annotated as mitochondrial are shown with blue dots. Well-known mtDNA-interacting proteins are shown with red dots and labelled with their gene names. Proteins located to the nucleus which were significantly increased after DNase treatment are shown with orange dots. **C**. Comigration and similar intensities of POLRMT and TFAM at ∼3 MDa in the control profile. **D**. POLG migrates at ∼1.9 MDa regardless addition of DNase. Segments of complexome profiles shown as heatmaps representing iBAQ values normalised to the maximum value within each pair of full profiles. Known interactors of POLG (pink) are shown in yellow, proteins comigrating with its ∼1.9 MDa fraction are shown in green. Each profile represents an average of three independent profiles.

To quantify these differences, we calculated Hausdorff effect sizes (*H_ES_*) using the COPAL tool (20). This parameter reflects how different the protein profiles are based on changes in both abundance and migration patterns. In several cases, however, the *H_ES_*values did not correlate with the observed changes due to the presence of multiple peaks or inconsistent identification of low abundant proteins. Thus, we also quantified the changes in intensity for individual peaks and tested if their appearance after DNase treatment was consistent. In parallel, we quantified the changes of total protein abundance between the triplicates using the total LFQ intensities of proteins in each profile (**Fig. 5B, Suppl. Data 5).** Only 74 out of 3800 detected proteins displayed significant changes upon DNase treatment, including mtDNA-interacting proteins TFAM, MTERF1, POLG2, UNG, and Twinkle. Other known mtDNA-binding proteins, such as EXOG, ENDOG, POLDIP2, were not significantly affected, in accordance to their transient interactions with mtDNA (41–43). Moreover, several nucDNA-interacting proteins such as histones, replication and repair factors, were enriched in the +DNase profile, further supporting the advantage of this approach. All these data showed the consistent improvement in detection of the mtDNA-binding proteins by CP after DNase treatment.

Interestingly, the mitochondrial RNA polymerase POLRMT (*M*_t_ 138.6 kDa) (44) migrated in the same *M*_a_ range as TFAM, peaking at ∼3.2 MDa, but it was not significantly affected by DNA digestion. The abundance of TFAM in this range in the untreated samples was similar to that of POLRMT (**Figure 5C**) and increased ∼5.5 times after DNase treatment. Considering that fully sized circular mtDNA and their bound components are expected not to enter the native gel in untreated samples, the ∼3.2 MDa peak of POLRMT and the DNase-insensitive portion of TFAM can represent the mtDNA-interacting transcription complex (45) that largely dissociates from mtDNA under regular solubilization conditions and therefore is detectable without the aid of the DNase treatment. The high *M*_a_ of the peak indicates the presence a nucleic acid component, possibly the nascent transcript. The other components of the transcription machinery TEFM and TFB2M displayed lower abundance and less defined profiles (**Figure 5A**), especially TFB2M, which might reflect their instability as parts of the complex, low cellular abundance or challenging MS identification of the peptides.

Notably, the occurrence of the ∼1.9 MDa fraction of POLG (*M*_t_ 140 kDa) did not change upon DNase treatment. This high *M*_a_ population of POLG has also been detected in a previous CP study (29), where it was suggested to be complexed with Twinkle and UNG. However, in our dataset, both of these proteins clearly migrated at a higher *M*_a_ than POLG and had much lower abundances (**Figure 5D**). Another suggested direct interactor of POLG, Top3α (46), was not present in that fraction either. In contrast, the proteins that displayed similar migration patterns in this mass range were the mitochondrial exonuclease G (EXOG), prohibitin, and the mitochondrial protein disaggregase CLPB. It has also been shown that POLG is degraded when it is not complexed with POLG2 (47). In this regard, the nature of this mtDNA- and POLG2-independent fraction of POLG deserves further attention.

Overall, we showed that the inclusion of DNase treatment in the standard CP workflow enhanced the detection of various (mt)DNA-binding proteins. Altogether, we established that hrCNE-based CP is a convenient approach for the investigation of mtRNA- and mtDNA-interacting proteins as well as nucleoprotein complexes. For all the following experiments, we set the hrCNE-based protocol with the DNase treatment step as the reference condition, as it enables consistent detection of the components of both mitochondrial DNA replication and gene expression machinery.

### Characterization of the high molecular mass range cluster of mtRNA- and mitoribosome-interacting proteins using RNase-treatment

In our improved complexome profiles, we observed multiple mtRNA-interacting proteins coinciding at ∼3 MDa (**Figure 2**). To better understand their relations and describe the behaviour of particular complexes in the native gel, we sought to determine their dependence on RNA. To this end, we generated a hrCNE-based profile using samples treated with both DNase I and RNase A (two replicates) and compared it with the control profile generated with only DNase-treated samples (three replicates) **(Suppl. Data 3)**. Bovine pancreas RNase A (RNase hereafter) cleaves both single- and double-stranded RNA, including the one found in RNA:DNA hybrids, thereby is expected to degrade all the accessible RNA in digitonin-solubilised mitochondrial samples.

In the resulting CP dataset, we selected a list of mitochondrial gene expression factors based on recent reviews (48–52) and arranged this list of 164 proteins according to their molecular functions and migration profiles **(Suppl. Data 3)**. Proteins involved in mt-mRNA processing (e.g. FASTKD family, GRSF1), stability (e.g. LRPPRC, SLIRP) and degradation (e.g. SUV3) as well as tRNA-related enzymes, predominantly migrated at lower *M*_a_. In contrast, mitoribosomes, mt-rRNA-processing enzymes, and some of the transcription factors were consistently detected at the megadalton mass range.

Most of the selected proteins were related to the mitoribosomes and mitochondrial translation. In total, 81 out of 82 structural mitoribosomal proteins (MRPs) were detected with only the smallest protein MRPL34 missing (53). In the control profile, the LSU migrated mainly at the slice corresponding to ∼3.2 MDa, while the SSU was found in the ∼2.3-3.2 MDa region (**Figure 6A**). In the RNase profile, each of the mitoribosomal subunits displayed two peaks: individual peaks at ∼1.5 MDa and ∼1.8 MDa for SSU and LSU, respectively, and a ∼2.8 MDa peak where both subunits overlapped. This indicated that the RNase treatment caused partial dissociation of the mitoribosome. Overall, the *M*_a_ of the mitoribosome-related peaks that appeared after RNase treatment roughly matched the calculated masses of the ribonucleoprotein complexes, i.e. 2.9 MDa for monosome, 1.8 MDa for LSU, and 1.1 MDa for SSU. In the control profile, the mitoribosomes migrated at a much higher *M*_a_, reaching almost 4 MDa. *In vivo*, the fully assembled mitoribosome, composed of the 82 MRPs, 12S (SSU) and 16S (LSU) mt-rRNAs, and a structural mt-tRNA^Val^ (LSU) (54,55), is associated with not only various auxiliary proteins, but also a translating mt-mRNA and the nascent polypeptide, of course resulting in a larger complex. Before the monosome forms, mitoribosomal subunits undergo individual assembly and maturation involving multiple intermediate states (56–59). These intermediates can have similar apparent masses due to presence or absence of just a few auxiliary factors. The assembly of LSU involves an especially large number of interactors (60). We thus suggest that the mitoribosome-related particles migrate in the hrCNE gel as a mixture of LSU and SSU intermediates and fully assembled mitoribosomes with preserved mt-rRNA and associated with mt-mRNA and interacting proteins, similar to what has been described for sucrose density gradients. Because of the low resolution of the upper part of the native gel and similar *M*_a_ of the complexes, it is currently difficult to dissect such potential subpopulations. However, for future research, this limitation might be circumvented by adjusting the polyacrylamide gradient and increasing the number of slices of this specific region of the gel, as described by others (61,62).

**Figure 6.**
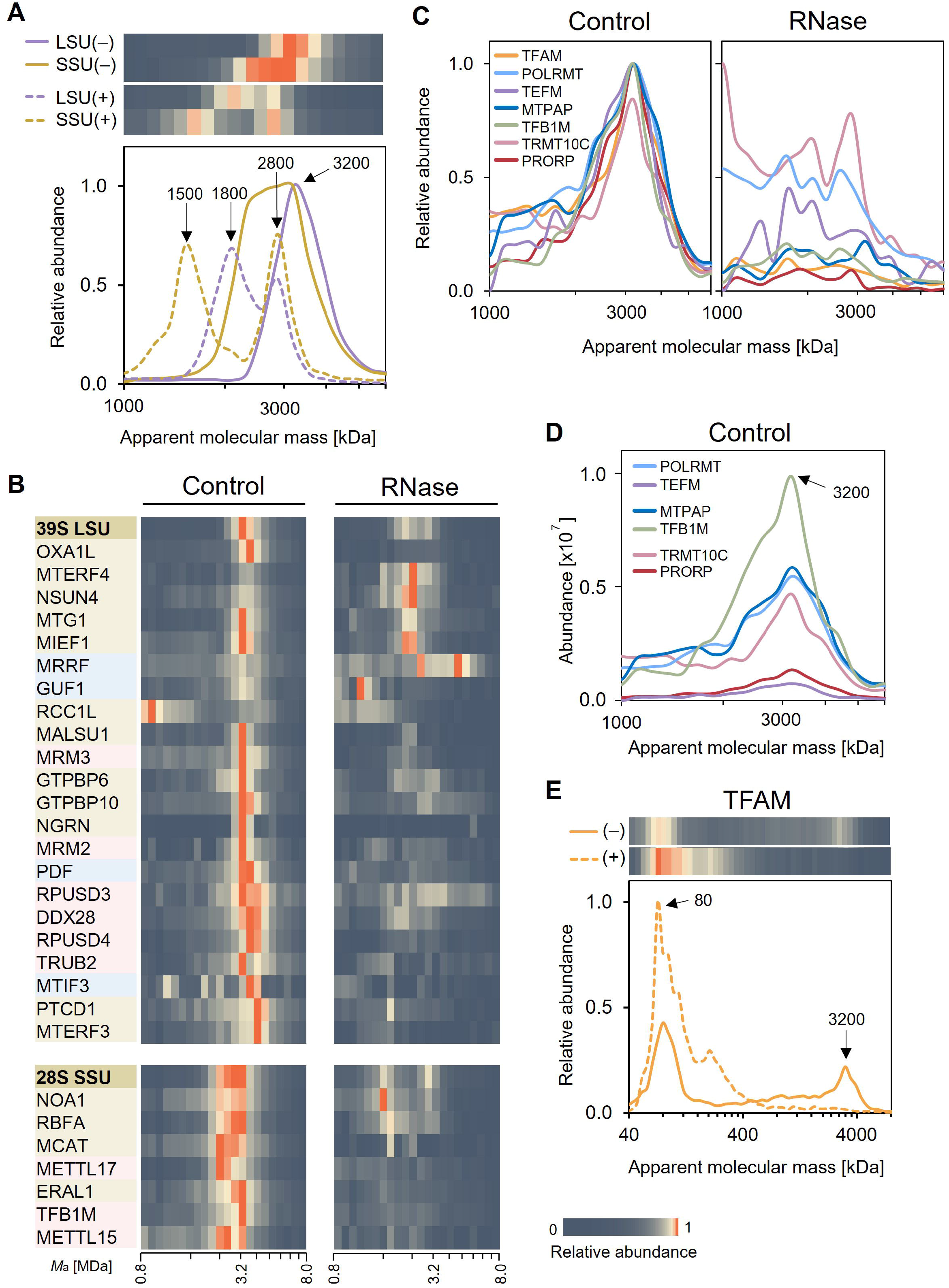
Abundance profiles of mitochondrial nucleoprotein complexes in the control and samples treated with RNase A (+RNase). Segments of complexome profiles are shown as heatmaps and line charts. The relative abundances were normalised to the maximum value within each pair of shown segments. The profiles represent an average of three control (-) or two RNase (+) independent profiles. Apparent molecular masses of the observed peaks are indicated with arrows in kDa. **A.** Profiles of mitoribosomal subunits LSU and SSU. **B**. Proteins involved in mitoribosome assembly (light brown), rRNA modification (pink) and mitochondrial translation (blue) comigrate with the respective mitoribosomal subunits (brown). Proteins are grouped based on similarity of migration patterns. **C**. Comigration of transcription and mtRNA-processing factors in control and +RNase profiles depicted by relative abundances. **D**. Abundances of the comigrating transcription and mtRNA-processing factors in the control profile. TEFM and PRORP have lower intensities compared to the other proteins. TFAM is not shown here due to a much higher abundance in the control profile. **E**. Differences in the migration profile of TFAM. The ∼3200 kDa peak of TFAM disappears after RNase treatment.

The proteins involved in mitochondrial translation, mt-rRNA processing and mitoribosome assembly, largely comigrated with the mitoribosomes in the 2.2-4.0 MDa range (**Figure 6B** – selected proteins, **Suppl. Data 3** – full list). We could clearly distinguish assembly factors of the SSU and many of the LSU, although some enzymes involved in 16S rRNA maturation (DDX28, RPUSD4, RPUSD3) or stability (PTCD1) showed variable patterns. After RNase treatment, some assembly factors remained attached to the distorted mitoribosomal subunits. For example, the LSU-binding complex MTERF4-NSUN4 as well as MALSU1 and MTG1 comigrated with the 1.8 MDa LSU peak, and the 12S rRNA chaperone ERAL1 comigrated with the 1.5 MDa SSU peak. GTPBP6 (LSU), RBFA and NOA1 (SSU) additionally comigrated with the residual monosome. Other proteins displayed very dispersed profiles upon RNase treatment, e.g. RPUSD4 and TRUB2 (LSU), and METTL15 (SSU). Interestingly, we also detected the malonyl-CoA-Acyl carrier protein transacylase (MCAT), a protein that has only recently been shown to be a SSU assembly factor (63), to comigrate with SSU in an RNA-dependent manner, proving the applicability of our method. We furthermore observed that some proteins without confirmed roles in mitochondrial gene expression displayed similar migration patterns and sensitivity to RNase as those from the known mitoribosome-interacting proteins, e.g. TOMM34, LETMD1, ACOT13, and SIRT4 (**Fig. S3**). This analysis confirmed the utility of hrCNE-based CP for analysis of mitoribosome interactors.

The other type of proteins found at the same mass range were mitochondrial transcription and mtRNA-processing factors (**Figure 6C**), namely, TFAM, transcription elongation factors POLRMT and TEFM, a small amount of the initiation factor TFB2M, mitochondrial poly(A)-polymerase MTPAP (64), 12S rRNA dimethyladenosine transferase TFB1M (65), and portions of the two RNase P subunits, TRMT10C and PROPR, involved in cleavage and maturation of mt-tRNAs (66). In the control, these proteins migrated in a range from ∼1 to ∼3.2 MDa, peaking at ∼3.2 MDa similarly to LSU. This migration pattern was distinguishable from that of LSU and most of its assembly factors as it was more spread towards lower *M*_a_ (see Suppl. Data 3), suggesting these proteins belong to a separate complex migrating in that mass range. After the RNase treatment, their signals largely overlapped and were characterised by scattered profiles between ∼0.5 and ∼3.2 MDa, indicating the presence of RNA partially protected from digestion by the interacting proteins.

It should be noted that the intensity of TEFM was 9 times lower compare to POLRMT, and the intensity of PROPR was 4.5 times lower than TRMT10C, while POLRMT, MTPAP, TRMT10C and TFB1M had similar intensities in the overlapping region (**Figure 6D**). The third subunit of RNase P, HSD17B10, partially comigrated with its interactors, although it had a very high abundance at lower *M*_a_. Such differences in the abundance among known protein interactors may indicate the presence of multiple complexes coinciding at similar *M*_a_ (e.g. the case of TFAM discussed in the previous section), a different stoichiometry of proteins in a given complex, or instability of some components during native electrophoresis. Despite this, the observed RNA-dependent comigration of the transcription factors with the mtRNA-processing factors was consistent and points towards a simultaneous interaction of all these proteins with the nascent transcript.

Curiously, TFAM displayed a very high sensitivity to RNase (**Figure 6E**), despite primarily binding mtDNA and not mtRNA (67). As we previously suggested, the majority of the high *M*_a_ fraction of TFAM is likely bound to the partially digested mtDNA, as its intensity increased after DNA digestion, with only a portion of TFAM belonging to the transcription complex. Thus, a possible explanation of the RNase sensitivity could be that removal of mtRNA made mtDNA more accessible for DNase digestion, eliminating the protective effect and therefore releasing TFAM.

It is important to note that besides the expected effects of the RNase treatment on RBPs, such as components of mitoribosomes, we observed some disturbance in CI, CV and the proteasome **(Suppl. Figure S2A)**. In fact, a previous study showed that some proteins aggregate and pellet from cell lysates when RNA is enzymatically digested, even if they lack RNA-binding motifs, among them the aforesaid protein complexes (68). However, the vast majority of the mitochondrial complexome was not significantly affected by this treatment, as exemplified in our selected reference complexes such as prohibitin, VDAC, CII, and OGDH **(Suppl. Figure S2B),** thus verifying the overall quality of the presented dataset.

### Investigating protein interactions of the mitochondrial gene expression system by hrCNE-based complexome profiling

The complexity of the mitochondrial gene expression system results from its many, often transient, protein-protein and nucleic acid-protein interactions. The experimental strategy developed here provides a reliable option for identifying protein interactor candidates and better comprehension of such a system. However, in the current setup, many proteins of different nature migrated at the similar mass range, such as mtDNA-, mtRNA- and mitoribosome-interacting proteins. It should be noted that similar mobility in native gels does not immediately mean that the identified proteins truly exist as a complex. Thus, prior experimental manipulation of the complexes under study in combination with CP can provide more information on the specific composition and formation of complexes. This has been applied for unravelling the assembly process of human CI where its assembly intermediates were tracked by complete inhibition of mitochondrial protein synthesis by chloramphenicol followed by release of the inhibition (69).

Here, to further demonstrate the utility of our optimised CP approach and to gain new insight about the interactions of the mitochondrial gene expression system, we similarly performed an inhibition-recovery experiment but adding ethidium bromide (EtBr) to the growth medium. This treatment leads to selective inhibition of both mitochondrial RNA and DNA synthesis in cultured cells (70) that can be reversed after removal of EtBr (71). Thus, short-term treatment with EtBr is expected to deplete mitochondrial DNA, rRNAs, tRNAs, and mRNAs with subsequent reduction of mitochondrial protein complexes containing mtDNA-encoded products, i.e. mitoribosomes, CI, CIII, CIV, and CV. After the removal of EtBr, mtDNA-/RNA synthesis resumes (72,73) and gradual restoration of the affected complexes is expected. These conditions thus allow us to study the formation processes of protein-nucleic acid complexes while they happen in genetically identical cells and before mitochondria undergo a large complexome remodeling upon prolonged mtDNA depletion (29).

To this end, we generated mitochondrial complexome profiles from HEK293 cells treated with a moderate concentration (30 ng/ml) of EtBr for three days (72 h) and from recovering cells of one day (24 h) after EtBr removal and compared them with the control dataset **(Suppl. Data 4)**.

### Mitochondrial gene expression rapidly recovers after removal of EtBr

We first evaluated the inhibitory effect of EtBr on mitochondrial gene expression and the level of its recovery upon EtBr removal in the samples used for CP. The analysis of the mtDNA relative content showed that after three days of EtBr-exposure the mtDNA levels decreased to ∼33% of the initial levels and then restored to ∼57% after one day without EtBr **(Figure S4A)**. The levels of both 12S and 16S mt-rRNAs were greatly reduced in response to the treatment and rapidly restored to almost the initial levels after the recovery phase as detected by Northern blotting **(Figure S4B)**. The twelve detected mtDNA-encoded OXPHOS proteins by CP were present at on average 23% of the control levels in the treated sample and showed a slow increase no more than 27.5% of control after recovery **(Figure S4D),** as estimated from their total abundances per complexome profile. Altogether, the EtBr-treated sample displayed reduced levels of mtDNA, mt-rRNA and mtDNA-encoded proteins and whereas the levels of mtDNA and mt-rRNA were rapidly restored after EtBr removal, this was not yet reflected by a full recovery of mitochondrial protein synthesis and consequently the OXPHOS complexes.

We then assessed the general changes of the mitochondrial proteome caused by EtBr and its subsequent removal (**Figure 7**). As anticipated, a dramatic reduction of assembled OXPHOS complexes I, III, IV, and V was detected upon EtBr treatment. Since CII is the only OXPHOS complex without mtDNA-encoded subunits, its levels were unsurprisingly not affected. The levels of mitoribosomes were greatly decreased as well (∼18% of control), reflecting the depletion of mt-rRNAs. After 24 h of recovery, the abundance of mitoribosomes increased to ∼61% of the control levels, whereas the affected OXPHOS complexes mostly remained unchanged. Since an efficient restart of the mitochondrial DNA replication and gene expression is anticipated, each pathway should be reactivated in a coordinated fashion. Accordingly, our findings indicate that both the mtDNA replication and assembly of mitoribosomes have a higher priority and occur faster during recovery from mtDNA-depletion compared to the protein synthesis using the remaining translation machinery. The latter possibly takes more time as additional molecular events should take place first; e.g. mt-mRNA and mt-tRNAs maturation, synthesis and import of nuclear-encoded OXPHOS subunits and assembly factors, etc.

**Figure 7.**
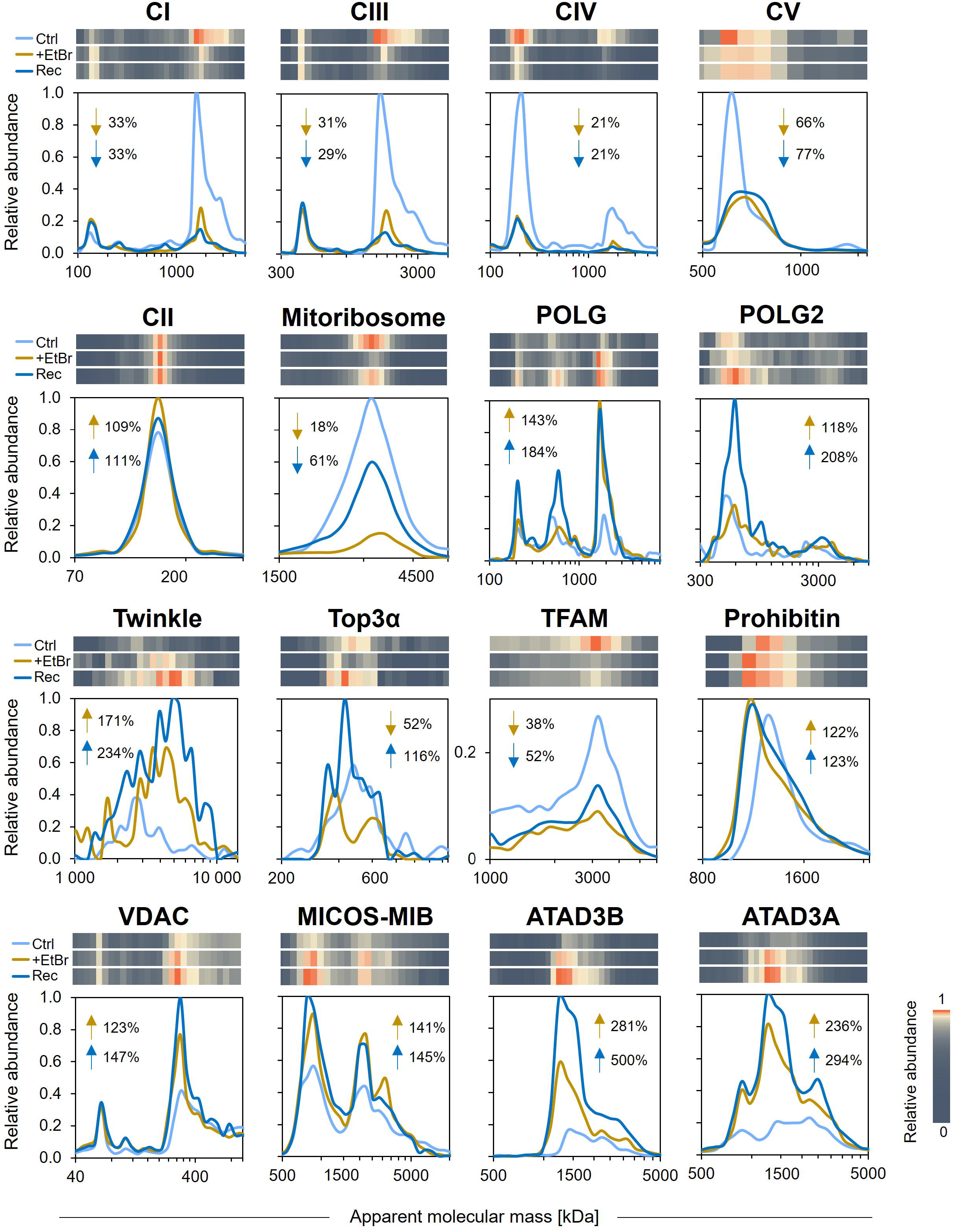
Comparison of the abundance profiles of mitochondrial proteins/complexes and mtDNA-binding proteins in the control, ethidium bromide (EtBr) treatment and recovery conditions. Segments of complexome profiles are shown as heatmaps and line charts. The relative abundances represent iBAQ values normalised to the maximum value within the three full profiles. The control (Ctrl) profile represents the average of three independent profiles. The EtBr treatment (+EtBr) and recovery (Rec) profiles represent single replicates. The differences in abundance between the conditions were determined using the summed iBAQ values across the shown mass ranges, compared and expressed as percentage of the control value. In each case, the resulting values for the treatment and recovery conditions are shown next to the profiles.

Furthermore, the mtDNA replication factors Polγ and Twinkle showed no reduction upon EtBr treatment and were highly enriched after recovery, together with Top3α. In contrast, TFAM followed the trend of mtDNA levels, as reflected by the intensity of its high *M*_a_ peak estimated at ∼38% of the control levels in the treatment and ∼52% in the recovery profiles. These observations suggest that mitochondria actively recover mtDNA content after the release of replication inhibition.

Finally, we detected elevated levels (up to 5-fold of control levels) of ATAD3A and ATAD3B proteins in both treatment and recovery conditions. While VDAC and MICOS-MIB complexes were slightly increased, the prohibitin complex displayed a downward mass shift without a change in intensity. These observations likely depict the reorganization of mitochondrial membrane architecture in response to EtBr-induced stress and mtDNA depletion (74,75).

### Mitoribosomal subassemblies accumulate at lower molecular mass range when mitochondrial gene expression is compromised

Since we observed a clear depletion-recovery pattern for the mitoribosome, we next wondered whether we can capture subassemblies of mitoribosomal subunits by CP. The mitoribosome assembly is a complex process that consists of mt-rRNA synthesis and its processing, including tRNA cleavage, rRNA modification and folding; sequential recruitment of assembly factors and MRPs to mt-rRNAs; and joining of the subunits into the 55S monosome (76). The assembly of SSU and LSU has been extensively studied in recent years by analysis of cryo-EM structures at atomic resolution, which detailed multiple assembly intermediates (56-58,63,77). In addition, the hierarchy of MRPs binding to mt-rRNAs was investigated in a pulse-chase stable isotope labeling by amino acids in cell culture (SILAC) study that described whether particular MRPs incorporated in the mitoribosomes at early or late stages of assembly (78).

Our hrCNE complexome profiles showed an accumulation of a number of MRPs in clusters below the main LSU and SSU populations which were different between the conditions (**Figure 8**). To understand the origin of these clusters, we mapped their identified components to the 3D structure of human mitoribosome (PDB 7QI4) and found that the comigrating proteins were generally located close to each other in the complex. We also compared the composition of the observed subassemblies with the kinetic model proposed by Bogenhagen et al. (78) and found that the majority of the proteins clustered in the SILAC study with some exceptions.

**Figure 8.**
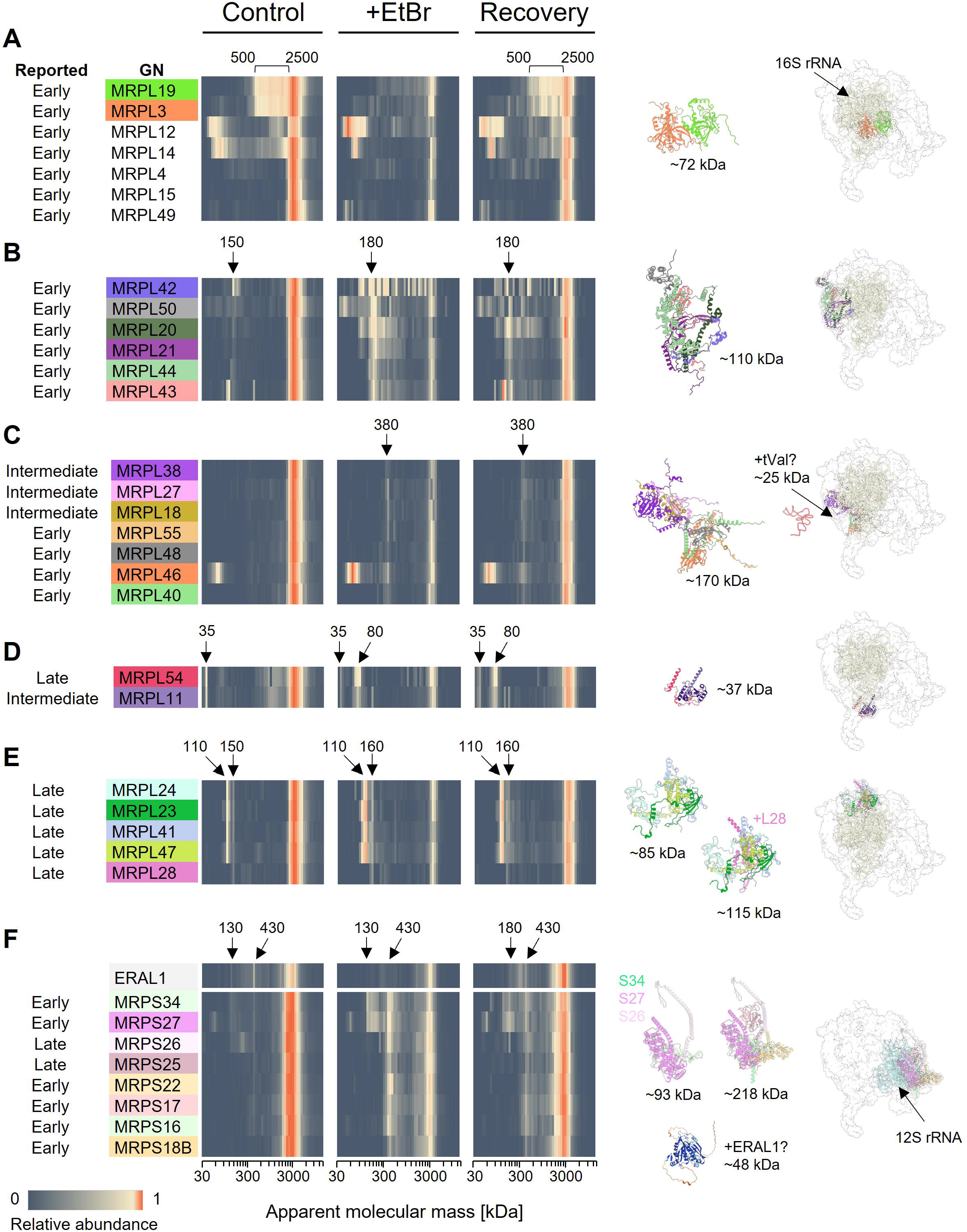
Comparison of the abundance profiles of mitoribosomal structural proteins in control, EtBr treatment and recovery conditions. A-F. Segments of complexome profiles are shown as heatmaps. The relative abundances represent iBAQ values normalised to the maximum value across the three full profiles. The control profile represents the average of three independent profiles. The treatment and recovery profiles represent single replicates. Apparent molecular masses of the observed clusters are indicated with arrows. The cartoons (right side) show the mitoribosomal proteins of the cluster and their position on the 3D structure of mitoribosome (PDB: 7QI4) relatively to mt-rRNA. The predicted structure of ERAL1 was retrieved from AlphaFold (105). The theoretical masses of the protein components/subassemblies are shown next to the structures. The proposed assembly stages in (78) are indicated for each protein. The proteins in the list are colored according to their respective color in the 3D structure.

The proteins reported to be among the first to bind 16S rRNA, namely MRPL3 and MRPL19, accumulated between 500 kDa and the main LSU peak (∼2500 kDa) (**Figure 8A**). The intensity of this population decreased after EtBr treatment and recuperated upon recovery, following the depletion-recovery course of mt-rRNA. Thus, these proteins likely indicate the presence of maturing LSU intermediates containing 16S mt-rRNA.

The early-stage proteins MRPL14 and MRPL12 showed a similar but less pronounced pattern, while the other early-stage MRPLs were mainly absent outside the LSU in the control but formed some lower molecular mass clusters in the treatment and recovery profiles. Specifically, six early-stage MRPLs comigrated at ∼180 kDa (**Figure 8B**). These proteins have been previously observed to interact in absence of mt-rRNA in rho0 cells, implying that they can form an subassembly without the rRNA scaffold (29). Three other early-stage MRPLs comigrated with four late-stage proteins at ∼380 kDa (**Figure 8C**). The total *M*_t_ of this cluster was 170 kDa, which was significantly lower than the observed *M*_a_, suggesting additional presence of other proteins or fragments of 16S rRNA (*M*_t_ ∼500 kDa) that might derive from mitoribosome disintegration induced by the aberrant assembly.

Interestingly, two peripheral 16S-binding proteins MRPL54 and MRPL11, which were reported to arrive at the mitoribosome at the later assembly stages, displayed a similar pattern to the early binding proteins as they accumulated below the LSU peak in the control and recovery samples (**Figure 8D**). In addition, they showed co-migration at 35 kDa and 80 kDa, although they were not reported to interact in a free form. 35 kDa would fit in the size of a MRPL54-MRPL11 heterodimer (*M*_t_ 37 kDa). Conversely, the 80 kDa peak, which was mainly present in the EtBr and recovery profiles, could represent the proteins with a fragment of mt-rRNA. These two proteins were highly sensitive to the RNase treatment after which they formed similar fragments at 90-120 kDa **(Suppl. Data 3)**.

Lastly, several late-stage MRPLs formed two clusters at 110 and 160 kDa that differed by the presence of MRPL28 (**Figure 8E**), which was not reported to cluster with these proteins in the SILAC study.

For the SSU, we observed a cluster of eight proteins (*M*_t_ 218 kDa) migrating at 430 kDa that was more pronounced in the treatment and recovery than in control (**Figure 8F**). The 12S mt-rRNA chaperone ERAL1 (*M*_t_ 48 kDa) comigrated with this cluster in the control profile and was detected in a similar mass range in the treatment and recovery ones. With the exception of MRPS25 and MRPS26, these proteins were reported as early-binding and to arrive to the mitoribosome at the same time. This cluster of MRPSs was also found to persist in rho0 cells at ∼400 kDa (29), suggesting that the above proteins can interact with each other in absence of mt-rRNA. The discrepant *M*_a_ of the cluster suggests the presence of other protein interactors. The dispersed pattern observed in the treatment and recovery profiles possibly suggests the simultaneous incorporation of these proteins to the assembling SSU.

Overall, we observed multiple MRPs comigrating at lower *M*_a_. While the majority of the clustered proteins were reported to interact in the SILAC study, we observed additional interactions. One explanation for the observed differences with the SILAC study is that CP directly identified mitoribosome subassemblies and their possible interactors. This can be contrasted and adds another layer of information compared to the SILAC based study by Bogenhagen et al. (78). There, the order of assembly was inferred from the timeline of arrival of proteins in gradient-purified (near) full-size mitoribosomes. This methodology thus excluded the detection of possible intermediates that were only a fraction of the size of mitoribosomes. Thus, while the suggested MRPL54-MRPL11 and other possible interactions might reflect early interactions in mitoribosome assembly, the arrival of these proteins in the almost fully mature mitoribosome would be interpreted as intermediate to late by the SILAC method.

### Several transcription, mtRNA-processing and early SSU assembly factors are largely resistant to EtBr treatment and comigrate

The mitoribosome assembly factors and mt-rRNA-interacting proteins generally followed the trend of depletion-recovery observed for SSU and LSU (**Figure 9A**). However, several proteins showed little or no depletion and only a moderate increase after removal of EtBr. Particularly, 12S mt-rRNA-binding assembly factors NOA1, ERAL1, TFB1M and MCAT, comigrated with SSU and did not change substantially their migration patterns and abundance in response to EtBr (67%, 55%, 62%, 67% of total abundance in control, respectively) (**Figure 9C**). These proteins are involved in maturation of the decoding center during early stages of SSU assembly together with METTL17 and RBFA (63). Conversely, these two proteins appeared more affected by EtBr (decreased by 5- and 4-fold, respectively) (**Figure 9C**). Since the abundance of the SSU in the +EtBr profile was still higher compared to these EtBr-“resistant” proteins (**Figure 9D**), they likely correspond to an SSU assembly intermediate that was not depleted by EtBr or was still able to assemble under these circumstances.

**Figure 9.**
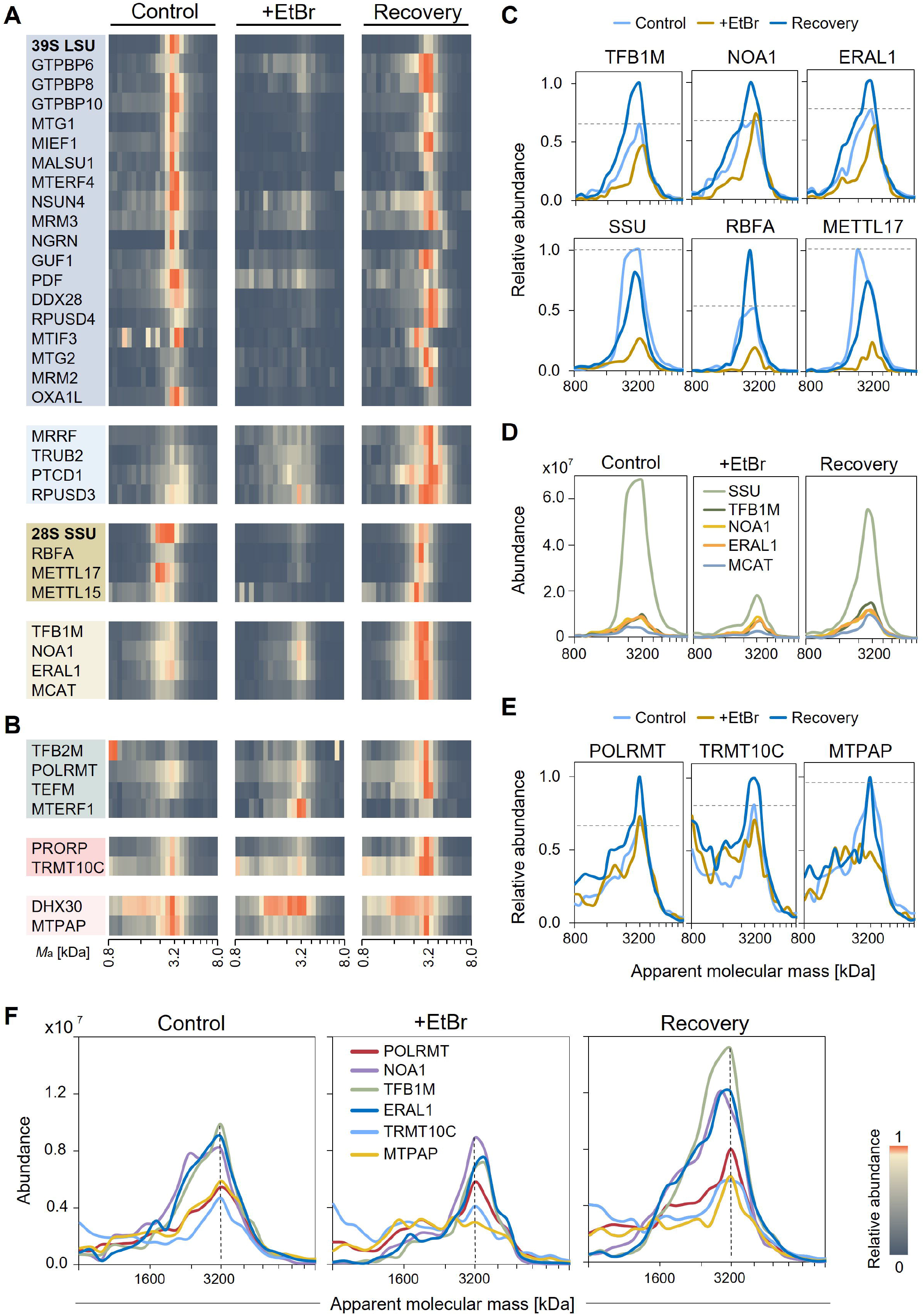
Comparison of the abundance profiles of mitoribosome assembly factors and mtRNA-binding proteins in control, EtBr treatment and recovery conditions. Segments of complexome profiles are shown as heatmaps and line charts. The relative abundances represent iBAQ values normalised to the maximum values within the three shown segments. The control profile represents an average of three independent profiles. The treatment and recovery profiles represent single replicates. A-B. The signals of the majority of assembly factors of LSU (blue) and SSU (brown) were depleted upon EtBr treatment and restored after the recovery phase. Some of the assembly factors were slightly depleted (light blue – LSU, light brown – SSU). Transcription factors (green) and two of the RNase P subunits (pink) were not depleted as well. mtRNA-binding proteins MTPAP and DHX30 (light pink) showed more dispersed patterns but were not depleted. C. Depletion and recovery trends of SSU and its selected assembly factors from the panel A. D. Abundances of SSU and its assembly factors TFB1M, NOA1, ERAL1 and MCAT. E. Depletion and recovery trends of the representative mtRNA-binding proteins from the panel B. The dashed lines indicate the maximal protein intensities in the control profile. F. Abundances of POLRMT, TRMT10C, MTPAP, TFB1M, NOA1 and ERAL1. The proteins overlapped at 3200 kDa or neighboring slices and had comparable intensities.

Next, we observed that transcription factors TFB2M, POLRMT, TEFM, comigrating at ∼3.2 MDa, were not depleted by EtBr either despite the demonstrated inhibition of mtRNA synthesis, and were increased in the recovery (**Figure 9B**). The transcription termination factor MTERF1 (**Figure 9B**), which was barely detected in control, was elevated in both treatment and recovery, and migrated at a similar *M*_a_. In addition, two subunits of RNase P, MTPAP and the poorly characterised mtRNA-helicase DHX30 (79) were only slightly affected by EtBr (**Figure 9B**). In the previous section, we observed RNase-sensitive comigration of the transcription and mtRNA-processing factors, suggesting they all simultaneously interact with nascent mtRNA. Generally, exposure to EtBr is thought to inhibit mitochondrial transcription, which is supported by the very low levels of mt-rRNAs detected after three days of the treatment, and data from similar studies (80–82). Nevertheless, the presence of residual transcription under EtBr treatment has previously been observed in different cell lines for both one-day treatments (83,84) and longer regimes (85,86). Taking this into consideration, we can imply that regardless the low mitochondrial transcript levels, these proteins can be a part of the mtRNA transcription-processing complex that, for example, shows extensive stalling.

Interestingly, these two groups of relatively unaffected by the EtBr treatment proteins _ the early-stage SSU assembly factors and mtRNA transcription and processing factors, showed very similar migration profiles in all tested conditions. The comigration was particularly clear for POLRMT, NOA1, TFB1M, ERAL1, TRMT10C and MTPAP (**Figure 9F**). This observation might indicate that the transcription and mtRNA-processing proteins interact with a subpopulation of SSU that contains the early assembly factors TFB1M, NOA1 and ERAL1. Since in human mitochondria transcription and translation are supposed to be spatially separated (87), this comigration unlikely represents a direct connection between transcription and translation machineries. Nevertheless, coupling mechanisms between transcription and RNA processing have been described for various species (88–90), including the direct interaction of some RNA processing factors with the RNA polymerase during transcription (91–93). An existence of such interaction in human mitochondria could explain these observations.

Importantly, the interaction between POLRMT, TFB1M and SSU has been previously demonstrated in immunoprecipitation experiments. Similarly to our observation, it was found to be persistent after a short EtBr treatment (3 h) (94). The authors proposed that POLRMT directly interacts with SSU and has a transcription-independent role in mitoribosome biogenesis, as they observed reduced 12S mt-rRNA methylation levels and less of fully assembled mitoribosomes when POLRMT was overexpressed. Here, we observed that SSU comigrated not only with POLRMT, but also with other transcription factors and mtRNA-processing factors, which were not tested in the aforementioned study. Taking into consideration the fact that EtBr treatment does not necessarily fully inhibit transcription initiation, mtRNA-protein interactions could persist after the treatment. Subsequently, the detected interaction of POLRMT and other transcription factors, mtRNA processing factors and a fraction of mitoribosomes might correspond to a mt-rRNA “processosome” that links transcription of 12S and 16S rRNAs, their processing, and the beginning of mitoribosome maturation.

In line with this, the human mitochondrial ribosome assembly has been previously suggested to start co-transcriptionally (5,78,95), similarly to the assembly of eukaryotic pre-ribosomes in the nucleolus (96) and assembly of bacterial ribosomes (97). This phenomenon has not yet been elucidated in detail for human mitochondria. It has been shown, however, that LSU can partially assemble in absence of mtRNA processing (95), suggesting that some MRPLs can bind to still transcribing mt-rRNA. A range of MRPs (5) and mitoribosome assembly factors have been found to interact with nucleoids, including the above-discussed SSU assembly factors NOA1 (98), ERAL1 and TFB1M (99), indicating they can bind nascent mt-rRNA. In yeast mitochondria, an existence of a closely interconnected network of proteins involved in transcription, mtRNA processing and assembly of the SSU has recently been proposed (100). These findings together with the here presented data point to the an existance of a coupling mechanism between mitochondrial transcription and mitoribosome assembly in human cells as well, which deserves further investigation.

## Conclusions and perspectives

Complexome profiling is nowadays a relatively well-established method that is most often used to investigate the composition, assembly and defects of OXPHOS complexes and other abundant mitochondrial proteins. We have here shown that it pays-off to further develop this approach, not only to expand its applicability on the analysis of whole mitochondrial proteomes, but also for enabling proper identification of DNA-/RNA-protein complexes. To achieve this, we included some small but crucial adjustments to the standard CP protocol. We first established that usage of high-resolution clear native electrophoresis prevents disruption of nucleic acid-protein interactions that are otherwise not stable when samples are separated by the commonly used blue native electrophoresis. Together with careful sample preparation, we also included a DNA digestion step in the protocol that further enhanced the identification of mtDNA-binding proteins. Our findings showed that the method is highly reproducible based on the similar detection of both mtDNA- and mtRNA-interacting proteins in different biological replicates, independent preparations and gel runs. Furthermore, a controlled and reversible manipulation of the mitochondrial gene expression system in combination with our method have resulted in identification of not only known factors involved in such pathways, but also a number of potential novel interactors with relevance to mitochondrial biology.

Overall, our results present a proof-of-concept to show that optimization of CP for the study of nucleic acid-protein interactions in mitochondria, combined with experimental manipulation of its gene expression system, adds a new tool that holds a great potential towards comprehensive analysis of the dynamics, composition and assembly of the complexes involved. Few limitations such as reliable detection of small and low abundant proteins, validation of identified interactors, accurate proteomic quantification and MS-time have yet to be tackled. Nevertheless, our method can be complemented with other advanced approaches, e.g., data-independent acquisition (DIA)-MS, crosslinking-MS, SILAC and tandem mass tags labeling. These tools should help overcome such limitations by providing specific information about protein structure, interaction sites, turnover and maturation, as well as multiplexing sample analysis and improving the MS running times, sensitivity and quantification.

On the experimental design side, a more inclusive time course of depletion-recovery as well as the analysis of systematic knockdown and knockout cell-lines for mitochondrial gene expression factors could provide a wealth of new information that can complement the analysis of complexes by other methods such as cryo-EM. As our findings also indicate that RNA may still be complexed with proteins during separation by hrCNE, potential recovery of such molecules may allow their further identification by, for example, RNA sequencing. This possibility has however to be proven feasible and experimentally adapted to avoid degradation and protein contamination. Our optimised protocol might also complement other approaches specifically designed for characterization of RNA-binding proteins based on UV-crosslinking followed by transcriptomic and proteomic analyses (101,102).

Even though our method has been developed to explore mitochondrial DNA-/RNA-interacting proteins, in principle it can be applied for characterization of any type of nucleic acid-protein complexes regardless their location within cells, as long as their sizes are not larger than the pore size limits of native gels. This method may also offer an unbiased, comprehensive and native structure-compatible option to explore the protein interaction partners of RNA groups whose roles are starting to emerge; e.g., non-coding RNAs (ncRNAs) (103). Finally, we invite the community to inspect the dataset generated by applying our optimised method since it is equally useful for general examination of protein complexes from mitochondria and their interactions.

## Data availability

The mass spectrometry proteomics data (raw files and search results) have been deposited to the ProteomeXchange Consortium via the PRIDE (104) partner repository with the dataset identifier PXD040103. The complexome profiling data have been deposited to the CEDAR database (26) with the accession number CRX42. The complexome profiling results for each experiment can be found in the supplementary files.

## Acknowledgements

We are grateful to Prof. Ulrich Brandt for helpful discussions and access to native electrophoresis and mass spectrometry. We also thank Dr. Joanna Rorbach for carefully reading our manuscript and valuable comments, Dr. Ilka Wittig for access to computer equipment, and Kimberley Bos for laboratory work assistance.

## Author contributions

AP, AC-O and JNS initially conceived and designed this study. Experimental work was carried out by AP and AC-O. AC-O and AP performed LC/MS-MS analysis. All authors were involved in data analysis, interpretation of the results and manuscript writing. The final draft of the manuscript has been approved by all authors.

## References

1. Nunnari, J. and Suomalainen, A. (2012) Mitochondria: in sickness and in health. Cell, 148, 1145–1159.

2. Rackham, O. and Filipovska, A. (2022) Organization and expression of the mammalian mitochondrial genome. Nat Rev Genet, 23, 606–623.

3. Garrido, N., Griparic, L., Jokitalo, E., Wartiovaara, J., van der Bliek, A.M. and Spelbrink, J.N. (2003) Composition and dynamics of human mitochondrial nucleoids. Mol Biol Cell, 14, 1583–1596.

4. Jourdain, A.A., Boehm, E., Maundrell, K. and Martinou, J.C. (2016) Mitochondrial RNA granules: Compartmentalizing mitochondrial gene expression. J Cell Biol, 212, 611–614.

5. Bogenhagen, D.F., Martin, D.W. and Koller, A. (2014) Initial steps in RNA processing and ribosome assembly occur at mitochondrial DNA nucleoids. Cell Metab, 19, 618–629.

6. Cabrera-Orefice, A., Potter, A., Evers, F., Hevler, J.F. and Guerrero-Castillo, S. (2022) Complexome Profiling-Exploring Mitochondrial Protein Complexes in Health and Disease. Front Cell Dev Biol, 9, 796128.

7. Heide, H., Bleier, L., Steger, M., Ackermann, J., Drose, S., Schwamb, B., Zornig, M., Reichert, A.S., Koch, I., Wittig, I. et al. (2012) Complexome profiling identifies TMEM126B as a component of the mitochondrial complex I assembly complex. Cell Metab, 16, 538–549.

8. Wessels, H.J., Vogel, R.O., Lightowlers, R.N., Spelbrink, J.N., Rodenburg, R.J., van den Heuvel, L.P., van Gool, A.J., Gloerich, J., Smeitink, J.A. and Nijtmans, L.G. (2013) Analysis of 953 human proteins from a mitochondrial HEK293 fraction by complexome profiling. PLoS One, 8, e68340.

9. Rugen, N., Straube, H., Franken, L.E., Braun, H.P. and Eubel, H. (2019) Complexome profiling reveals association of PPR proteins with ribosomes in the mitochondria of plants. Molecular and Cellular Proteomics, 18, 1345–1362.

10. Chatzispyrou, I.A., Alders, M., Guerrero-Castillo, S., Zapata Perez, R., Haagmans, M.A., Mouchiroud, L., Koster, J., Ofman, R., Baas, F., Waterham, H.R. et al. (2017) A homozygous missense mutation in ERAL1, encoding a mitochondrial rRNA chaperone, causes Perrault syndrome. Human molecular genetics, 26, 2541–2550.

11. Cunatova, K., Reguera, D.P., Vrbacky, M., Fernandez-Vizarra, E., Ding, S., Fearnley, I.M., Zeviani, M., Houstek, J., Mracek, T. and Pecina, P. (2021) Loss of COX4I1 Leads to Combined Respiratory Chain Deficiency and Impaired Mitochondrial Protein Synthesis. Cells, 10.

12. Gardeitchik, T., Mohamed, M., Ruzzenente, B., Karall, D., Guerrero-Castillo, S., Dalloyaux, D., van den Brand, M., van Kraaij, S., van Asbeck, E., Assouline, Z., et al. (2018) Bi-allelic Mutations in the Mitochondrial Ribosomal Protein MRPS2 Cause Sensorineural Hearing Loss, Hypoglycemia, and Multiple OXPHOS Complex Deficiencies. Am J Hum Genet, 102, 685–695.

13. Palenikova, P., Harbour, M.E., Ding, S., Fearnley, I.M., Van Haute, L., Rorbach, J., Scavetta, R., Minczuk, M. and Rebelo-Guiomar, P. (2021) Quantitative density gradient analysis by mass spectrometry (qDGMS) and complexome profiling analysis (ComPrAn) R package for the study of macromolecular complexes. Biochim Biophys Acta Bioenerg, 1862, 148399.

14. Van Haute, L., Hendrick, A.G., D’Souza, A.R., Powell, C.A., Rebelo-Guiomar, P., Harbour, M.E., Ding, S., Fearnley, I.M., Andrews, B. and Minczuk, M. (2019) METTL15 introduces N4-methylcytidine into human mitochondrial 12S rRNA and is required for mitoribosome biogenesis. Nucleic Acids Res, 47, 10267–10281.

15. Munawar, N., Olivero, G., Jerman, E., Doyle, B., Streubel, G., Wynne, K., Bracken, A. and Cagney, G. (2015) Native gel analysis of macromolecular protein complexes in cultured mammalian cells. Proteomics, 15, 3603–3612.

16. Hensen, F., Potter, A., van Esveld, S.L., Tarrés-Solé, A., Chakraborty, A., Solà, M. and Spelbrink, J.N. (2019) Mitochondrial RNA granules are critically dependent on mtDNA replication factors Twinkle and mtSSB. Nucleic Acids Research, 47, 3680–3698.

17. Wittig, I., Braun, H.P. and Schagger, H. (2006) Blue native PAGE. Nat Protoc, 1, 418–428.

18. Wittig, I., Karas, M. and Schagger, H. (2007) High resolution clear native electrophoresis for in-gel functional assays and fluorescence studies of membrane protein complexes. Mol Cell Proteomics, 6, 1215–1225.

19. Tyanova, S., Temu, T. and Cox, J. (2016) The MaxQuant computational platform for mass spectrometry-based shotgun proteomics. Nat Protoc, 11, 2301–2319.

20. Van Strien, J., Guerrero-Castillo, S., Chatzispyrou, I.A., Houtkooper, R.H., Brandt, U. and Huynen, M.A. (2019) COmplexome Profiling ALignment (COPAL) reveals remodeling of mitochondrial protein complexes in Barth syndrome. Bioinformatics, 35, 3083–3091.

21. Rath, S., Sharma, R., Gupta, R., Ast, T., Chan, C., Durham, T.J., Goodman, R.P., Grabarek, Z., Haas, M.E., Hung, W.H.W. et al. (2021) MitoCarta3.0: an updated mitochondrial proteome now with sub-organelle localization and pathway annotations. Nucleic Acids Res, 49, D1541–d1547.

22. Marchiano, F., Haering, M. and Habermann, B.H. (2022) The mitoXplorer 2.0 update: integrating and interpreting mitochondrial expression dynamics within a cellular context. Nucleic Acids Res, 50, W490–499.

23. Smith, A.C. and Robinson, A.J. (2018) MitoMiner v4.0: an updated database of mitochondrial localization evidence, phenotypes and diseases. Nucleic Acids Research, 47, D1225–D1228.

24. Wittig, I., Beckhaus, T., Wumaier, Z., Karas, M. and Schagger, H. (2010) Mass estimation of native proteins by blue native electrophoresis: principles and practical hints. Mol Cell Proteomics, 9, 2149–2161.

25. de Hoon, M.J., Imoto, S., Nolan, J. and Miyano, S. (2004) Open source clustering software. Bioinformatics, 20, 1453–1454.

26. van Strien, J., Haupt, A., Schulte, U., Braun, H.P., Cabrera-Orefice, A., Choudhary, J.S., Evers, F., Fernandez-Vizarra, E., Guerrero-Castillo, S., Kooij, T.W.A. et al. (2021) CEDAR, an online resource for the reporting and exploration of complexome profiling data. Biochim Biophys Acta Bioenerg, 1862, 148411.

27. Alston, C.L., Veling, M.T., Heidler, J., Taylor, L.S., Alaimo, J.T., Sung, A.Y., He, L., Hopton, S., Broomfield, A., Pavaine, J. et al. (2020) Pathogenic Bi-allelic Mutations in NDUFAF8 Cause Leigh Syndrome with an Isolated Complex I Deficiency. Am J Hum Genet, 106, 92–101.

28. Salscheider, S.L., Gerlich, S., Cabrera-Orefice, A., Peker, E., Rothemann, R.A., Murschall, L.M., Finger, Y., Szczepanowska, K., Ahmadi, Z.A., Guerrero-Castillo, S. et al. (2022) AIFM1 is a component of the mitochondrial disulfide relay that drives complex I assembly through efficient import of NDUFS5. EMBO J, 41, e110784.

29. Guerrero-Castillo, S., van Strien, J., Brandt, U. and Arnold, S. (2021) Ablation of mitochondrial DNA results in widespread remodeling of the mitochondrial complexome. *EMBO J*, e108648.

30. Gorka, M., Swart, C., Siemiatkowska, B., Martinez-Jaime, S., Skirycz, A., Streb, S. and Graf, A. (2019) Protein Complex Identification and quantitative complexome by CN-PAGE. Sci Rep, 9, 11523.

31. Wortmann, S.B., Ziętkiewicz, S., Guerrero-Castillo, S., Feichtinger, R.G., Wagner, M., Russell, J., Ellaway, C., Mróz, D., Wyszkowski, H., Weis, D. et al. (2021) Neutropenia and intellectual disability are hallmarks of biallelic and de novo CLPB deficiency. Genet Med.

32. Sasarman, F., Brunel-Guitton, C., Antonicka, H., Wai, T., Shoubridge, E.A. and Consortium, L. (2010) LRPPRC and SLIRP interact in a ribonucleoprotein complex that regulates posttranscriptional gene expression in mitochondria. Mol Biol Cell, 21, 1315–1323.

33. Hevler, J.F., Albanese, P., Cabrera-Orefice, A., Potter, A., Jankevics, A., Misic, J., Scheltema, R.A., Brandt, U., Arnold, S. and Heck, A.J.R. (2023) MRPS36 provides a structural link in the eukaryotic 2-oxoglutarate dehydrogenase complex. Open Biol, 13, 220363.

34. Arguello, T., Peralta, S., Antonicka, H., Gaidosh, G., Diaz, F., Tu, Y.T., Garcia, S., Shiekhattar, R., Barrientos, A. and Moraes, C.T. (2021) ATAD3A has a scaffolding role regulating mitochondria inner membrane structure and protein assembly. Cell Rep, 37, 110139.

35. Merle, N., Feraud, O., Gilquin, B., Hubstenberger, A., Kieffer-Jacquinot, S., Assard, N., Bennaceur-Griscelli, A., Honnorat, J. and Baudier, J. (2012) ATAD3B is a human embryonic stem cell specific mitochondrial protein, re-expressed in cancer cells, that functions as dominant negative for the ubiquitous ATAD3A. Mitochondrion, 12, 441–448.

36. Yu, B., Pettitt, B.M. and Iwahara, J. (2020) Dynamics of Ionic Interactions at Protein-Nucleic Acid Interfaces. Acc Chem Res, 53, 1802–1810.

37. Alam, T.I., Kanki, T., Muta, T., Ukaji, K., Abe, Y., Nakayama, H., Takio, K., Hamasaki, N. and Kang, D. (2003) Human mitochondrial DNA is packaged with TFAM. Nucleic Acids Res, 31, 1640–1645.

38. Kaguni, L.S. (2004) DNA polymerase gamma, the mitochondrial replicase. Annu Rev Biochem, 73, 293–320.

39. Lee, Y.S., Kennedy, W.D. and Yin, Y.W. (2009) Structural insight into processive human mitochondrial DNA synthesis and disease-related polymerase mutations. Cell, 139, 312–324.

40. Gustafsson, C.M., Falkenberg, M. and Larsson, N.G. (2016) Maintenance and Expression of Mammalian Mitochondrial DNA. Annu Rev Biochem, 85, 133–160.

41. Cheng, X., Kanki, T., Fukuoh, A., Ohgaki, K., Takeya, R., Aoki, Y., Hamasaki, N. and Kang, D. (2005) PDIP38 associates with proteins constituting the mitochondrial DNA nucleoid. J Biochem, 138, 673–678.

42. Wiehe, R.S., Gole, B., Chatre, L., Walther, P., Calzia, E., Ricchetti, M. and Wiesmuller, L. (2018) Endonuclease G promotes mitochondrial genome cleavage and replication. Oncotarget, 9, 18309–18326.

43. Szymanski, M.R., Karlowicz, A., Herrmann, G.K., Cen, Y. and Yin, Y.W. (2022) Human EXOG Possesses Strong AP Hydrolysis Activity: Implication on Mitochondrial DNA Base Excision Repair. J Am Chem Soc, 144, 23543–23550.

44. Ringel, R., Sologub, M., Morozov, Y.I., Litonin, D., Cramer, P. and Temiakov, D. (2011) Structure of human mitochondrial RNA polymerase. Nature, 478, 269–273.

45. Hillen, H.S., Temiakov, D. and Cramer, P. (2018) Structural basis of mitochondrial transcription. Nat Struct Mol Biol, 25, 754–765.

46. Hangas, A., Kekäläinen, N.J., Potter, A., Michell, C., Aho, K.J., Rutanen, C., Spelbrink, Johannes N., Pohjoismäki, Jaakko L. and Goffart, S. (2022) Top3α is the replicative topoisomerase in mitochondrial DNA replication. Nucleic Acids Research, 50, 8733–8748.

47. Silva-Pinheiro, P., Pardo-Hernandez, C., Reyes, A., Tilokani, L., Mishra, A., Cerutti, R., Li, S., Rozsivalova, D.H., Valenzuela, S., Dogan, S.A. et al. (2021) DNA polymerase gamma mutations that impair holoenzyme stability cause catalytic subunit depletion. Nucleic Acids Res, 49, 5230–5248.

48. Hilander, T., Jackson, C.B., Robciuc, M., Bashir, T. and Zhao, H. (2021) The roles of assembly factors in mammalian mitoribosome biogenesis. Mitochondrion, 60, 70–84.

49. Jedynak-Slyvka, M., Jabczynska, A. and Szczesny, R.J. (2021) Human Mitochondrial RNA Processing and Modifications: Overview. Int J Mol Sci, 22.

50. Lavdovskaia, E., Hillen, H.S. and Richter-Dennerlein, R. (2022) Hierarchical folding of the catalytic core during mitochondrial ribosome biogenesis. Trends Cell Biol, 32, 182–185.

51. Miranda, M., Bonekamp, N.A. and Kuhl, I. (2022) Starting the engine of the powerhouse: mitochondrial transcription and beyond. Biol Chem, 403, 779–805.

52. Wang, F., Zhang, D., Zhang, D., Li, P. and Gao, Y. (2021) Mitochondrial Protein Translation: Emerging Roles and Clinical Significance in Disease. Front Cell Dev Biol, 9, 675465.

53. De Silva, D., Tu, Y.T., Amunts, A., Fontanesi, F. and Barrientos, A. (2015) Mitochondrial ribosome assembly in health and disease. Cell Cycle, 14, 2226–2250.

54. Amunts, A., Brown, A., Toots, J., Scheres, S.H.W. and Ramakrishnan, V. (2015) Ribosome. The structure of the human mitochondrial ribosome. Science, 348, 95–98.

55. Greber, B.J., Bieri, P., Leibundgut, M., Leitner, A., Aebersold, R., Boehringer, D. and Ban, N. (2015) Ribosome. The complete structure of the 55S mammalian mitochondrial ribosome. Science, 348, 303–308.

56. Cheng, J., Berninghausen, O. and Beckmann, R. (2021) A distinct assembly pathway of the human 39S late pre-mitoribosome. Nat Commun, 12, 4544.

57. Itoh, Y., Khawaja, A., Laptev, I., Cipullo, M., Atanassov, I., Sergiev, P., Rorbach, J. and Amunts, A. (2022) Mechanism of mitoribosomal small subunit biogenesis and preinitiation. Nature, 606, 603–608.

58. Lenarcic, T., Niemann, M., Ramrath, D.J.F., Calderaro, S., Flugel, T., Saurer, M., Leibundgut, M., Boehringer, D., Prange, C., Horn, E.K. et al. (2022) Mitoribosomal small subunit maturation involves formation of initiation-like complexes. Proc Natl Acad Sci U S A, 119.

59. Rebelo-Guiomar, P., Pellegrino, S., Dent, K.C., Sas-Chen, A., Miller-Fleming, L., Garone, C., Van Haute, L., Rogan, J.F., Dinan, A., Firth, A.E. et al. (2022) A late-stage assembly checkpoint of the human mitochondrial ribosome large subunit. Nat Commun, 13, 929.

60. Chandrasekaran, V., Desai, N., Burton, N.O., Yang, H., Price, J., Miska, E.A. and Ramakrishnan, V. (2021) Visualizing formation of the active site in the mitochondrial ribosome. Elife, 10.

61. Muller, C.S., Bildl, W., Klugbauer, N., Haupt, A., Fakler, B. and Schulte, U. (2019) High-Resolution Complexome Profiling by Cryoslicing BN-MS Analysis. J Vis Exp.

62. Schulte, U., den Brave, F., Haupt, A., Gupta, A., Song, J., Muller, C.S., Engelke, J., Mishra, S., Martensson, C., Ellenrieder, L., et al. (2023) Mitochondrial complexome reveals quality-control pathways of protein import. Nature, 614, 153–159.

63. Harper, N.J., Burnside, C. and Klinge, S. (2022) Principles of mitoribosomal small subunit assembly in eukaryotes. Nature.

64. Bratic, A., Clemente, P., Calvo-Garrido, J., Maffezzini, C., Felser, A., Wibom, R., Wedell, A., Freyer, C. and Wredenberg, A. (2016) Mitochondrial Polyadenylation Is a One-Step Process Required for mRNA Integrity and tRNA Maturation. PLoS Genet, 12, e1006028.

65. Liu, X., Shen, S., Wu, P., Li, F., Liu, X., Wang, C., Gong, Q., Wu, J., Yao, X., Zhang, H. et al. (2019) Structural insights into dimethylation of 12S rRNA by TFB1M: indispensable role in translation of mitochondrial genes and mitochondrial function. Nucleic Acids Res, 47, 7648–7665.

66. Bhatta, A., Dienemann, C., Cramer, P. and Hillen, H.S. (2021) Structural basis of RNA processing by human mitochondrial RNase P. Nat Struct Mol Biol, 28, 713–723.

67. Brown, T.A., Tkachuk, A.N. and Clayton, D.A. (2015) Mitochondrial Transcription Factor A (TFAM) Binds to RNA Containing 4-Way Junctions and Mitochondrial tRNA. PLoS One, 10, e0142436.

68. Aarum, J., Cabrera, C.P., Jones, T.A., Rajendran, S., Adiutori, R., Giovannoni, G., Barnes, M.R., Malaspina, A. and Sheer, D. (2020) Enzymatic degradation of RNA causes widespread protein aggregation in cell and tissue lysates. EMBO reports, 21.

69. Guerrero-Castillo, S., Baertling, F., Kownatzki, D., Wessels, H.J., Arnold, S., Brandt, U. and Nijtmans, L. (2017) The Assembly Pathway of Mitochondrial Respiratory Chain Complex I. Cell Metab, 25, 128–139.

70. Coppey-Moisan, M., Brunet, A.C., Morais, R. and Coppey, J. (1996) Dynamical change of mitochondrial DNA induced in the living cell by perturbing the electrochemical gradient. Biophys J, 71, 2319–2328.

71. Hayakawa, T., Noda, M., Yasuda, K., Yorifuji, H., Taniguchi, S., Miwa, I., Sakura, H., Terauchi, Y., Hayashi, J., Sharp, G.W. et al. (1998) Ethidium bromide-induced inhibition of mitochondrial gene transcription suppresses glucose-stimulated insulin release in the mouse pancreatic beta-cell line betaHC9. J Biol Chem, 273, 20300–20307.

72. Seidel-Rogol, B.L. and Shadel, G.S. (2002) Modulation of mitochondrial transcription in response to mtDNA depletion and repletion in HeLa cells. Nucleic Acids Res, 30, 1929–1934.

73. Alán, L., Špaček, T., Pajuelo Reguera, D., Jabůrek, M. and Ježek, P. (2016) Mitochondrial nucleoid clusters protect newly synthesized mtDNA during Doxorubicin- and Ethidium Bromide-induced mitochondrial stress. Toxicol Appl Pharmacol, 302, 31–40.

74. Sen, A., Kallabis, S., Gaedke, F., Jungst, C., Boix, J., Nuchel, J., Maliphol, K., Hofmann, J., Schauss, A.C., Kruger, M. et al. (2022) Mitochondrial membrane proteins and VPS35 orchestrate selective removal of mtDNA. Nat Commun, 13, 6704.

75. He, B., Yu, H., Liu, S., Wan, H., Fu, S., Liu, S., Yang, J., Zhang, Z., Huang, H., Li, Q. et al. (2022) Mitochondrial cristae architecture protects against mtDNA release and inflammation. Cell Rep, 41, 111774.

76. Lopez Sanchez, M.I.G., Kruger, A., Shiriaev, D.I., Liu, Y. and Rorbach, J. (2021) Human Mitoribosome Biogenesis and Its Emerging Links to Disease. Int J Mol Sci, 22.

77. Cipullo, M., Gese, G.V., Khawaja, A., Hallberg, B.M. and Rorbach, J. (2021) Structural basis for late maturation steps of the human mitoribosomal large subunit. Nat Commun, 12, 3673.

78. Bogenhagen, D.F., Ostermeyer-Fay, A.G., Haley, J.D. and Garcia-Diaz, M. (2018) Kinetics and Mechanism of Mammalian Mitochondrial Ribosome Assembly. Cell Rep, 22, 1935–1944.

79. Antonicka, H. and Shoubridge, E.A. (2015) Mitochondrial RNA Granules Are Centers for Posttranscriptional RNA Processing and Ribosome Biogenesis. Cell Rep, 10, 920–932.

80. Arnold, J.J., Sharma, S.D., Feng, J.Y., Ray, A.S., Smidansky, E.D., Kireeva, M.L., Cho, A., Perry, J., Vela, J.E., Park, Y. et al. (2012) Sensitivity of mitochondrial transcription and resistance of RNA polymerase II dependent nuclear transcription to antiviral ribonucleosides. PLoS Pathog, 8, e1003030.

81. Piechota, J., Szczesny, R., Wolanin, K., Chlebowski, A. and Bartnik, E. (2006) Nuclear and mitochondrial genome responses in HeLa cells treated with inhibitors of mitochondrial DNA expression. Acta Biochim Pol, 53, 485–495.

82. Lee, W., Choi, H.I., Kim, M.J. and Park, S.Y. (2008) Depletion of mitochondrial DNA up-regulates the expression of MDR1 gene via an increase in mRNA stability. Exp Mol Med, 40, 109–117.

83. Kotrys, A.V., Cysewski, D., Czarnomska, S.D., Pietras, Z., Borowski, L.S., Dziembowski, A. and Szczesny, R.J. (2019) Quantitative proteomics revealed C6orf203/MTRES1 as a factor preventing stress-induced transcription deficiency in human mitochondria. Nucleic Acids Res, 47, 7502–7517.

84. Falabella, M., Kolesar, J.E., Wallace, C., de Jesus, D., Sun, L., Taguchi, Y.V., Wang, C., Wang, T., Xiang, I.M., Alder, J.K. et al. (2019) G-quadruplex dynamics contribute to regulation of mitochondrial gene expression. Sci Rep, 9, 5605.

85. Warren, E.B., Aicher, A.E., Fessel, J.P. and Konradi, C. (2017) Mitochondrial DNA depletion by ethidium bromide decreases neuronal mitochondrial creatine kinase: Implications for striatal energy metabolism. PLoS One, 12, e0190456.

86. Chatre, L. and Ricchetti, M. (2013) Large heterogeneity of mitochondrial DNA transcription and initiation of replication exposed by single-cell imaging. J Cell Sci, 126, 914–926.

87. Zorkau, M., Albus, C.A., Berlinguer-Palmini, R., Chrzanowska-Lightowlers, Z.M.A. and Lightowlers, R.N. (2021) High-resolution imaging reveals compartmentalization of mitochondrial protein synthesis in cultured human cells. Proc Natl Acad Sci U S A, 118.

88. Geisberg, J.V., Moqtaderi, Z., Fong, N., Erickson, B., Bentley, D.L. and Struhl, K. (2022) Nucleotide-level linkage of transcriptional elongation and polyadenylation. Elife, 11.

89. Peck, S.A., Hughes, K.D., Victorino, J.F. and Mosley, A.L. (2019) Writing a wrong: Coupled RNA polymerase II transcription and RNA quality control. Wiley Interdiscip Rev RNA, 10, e1529.

90. Proudfoot, N.J., Furger, A. and Dye, M.J. (2002) Integrating mRNA processing with transcription. Cell, 108, 501–512.

91. Rodeheffer, M.S. and Shadel, G.S. (2003) Multiple interactions involving the amino-terminal domain of yeast mtRNA polymerase determine the efficiency of mitochondrial protein synthesis. J Biol Chem, 278, 18695–18701.

92. Glover-Cutter, K., Kim, S., Espinosa, J. and Bentley, D.L. (2008) RNA polymerase II pauses and associates with pre-mRNA processing factors at both ends of genes. Nat Struct Mol Biol, 15, 71–78.

93. Andrulis, E.D., Werner, J., Nazarian, A., Erdjument-Bromage, H., Tempst, P. and Lis, J.T. (2002) The RNA processing exosome is linked to elongating RNA polymerase II in Drosophila. Nature, 420, 837–841.

94. Surovtseva, Y.V. and Shadel, G.S. (2013) Transcription-independent role for human mitochondrial RNA polymerase in mitochondrial ribosome biogenesis. Nucleic Acids Res, 41, 2479–2488.

95. Rackham, O., Busch, J.D., Matic, S., Siira, S.J., Kuznetsova, I., Atanassov, I., Ermer, J.A., Shearwood, A.M., Richman, T.R., Stewart, J.B. et al. (2016) Hierarchical RNA Processing Is Required for Mitochondrial Ribosome Assembly. Cell Rep, 16, 1874–1890.

96. Turowski, T.W. and Tollervey, D. (2015) Cotranscriptional events in eukaryotic ribosome synthesis. Wiley Interdiscip Rev RNA, 6, 129–139.

97. Shajani, Z., Sykes, M.T. and Williamson, J.R. (2011) Assembly of bacterial ribosomes. Annu Rev Biochem, 80, 501–526.

98. He, J., Cooper, H.M., Reyes, A., Di Re, M., Kazak, L., Wood, S.R., Mao, C.C., Fearnley, I.M., Walker, J.E. and Holt, I.J. (2012) Human C4orf14 interacts with the mitochondrial nucleoid and is involved in the biogenesis of the small mitochondrial ribosomal subunit. Nucleic Acids Res, 40, 6097–6108.

99. Bogenhagen, D.F., Rousseau, D. and Burke, S. (2008) The layered structure of human mitochondrial DNA nucleoids. J Biol Chem, 283, 3665–3675.

100. Singh, A.P., Salvatori, R., Aftab, W., Aufschnaiter, A., Carlstrom, A., Forne, I., Imhof, A. and Ott, M. (2020) Molecular Connectivity of Mitochondrial Gene Expression and OXPHOS Biogenesis. Mol Cell, 79, 1051–1065 e1010.

101. van Esveld, S.L. and Spelbrink, J.N. (2021) RNA Crosslinking to Analyze the Mitochondrial RNA-Binding Proteome. Methods Mol Biol, 2192, 147–158.

102. Trendel, J., Schwarzl, T., Horos, R., Prakash, A., Bateman, A., Hentze, M.W. and Krijgsveld, J. (2019) The Human RNA-Binding Proteome and Its Dynamics during Translational Arrest. Cell, 176, 391–403 e319.

103. Rinn, J.L. and Chang, H.Y. (2012) Genome regulation by long noncoding RNAs. Annu Rev Biochem, 81, 145–166.

104. Perez-Riverol, Y., Bai, J., Bandla, C., Garcia-Seisdedos, D., Hewapathirana, S., Kamatchinathan, S., Kundu, D.J., Prakash, A., Frericks-Zipper, A., Eisenacher, M. et al. (2022) The PRIDE database resources in 2022: a hub for mass spectrometry-based proteomics evidences. Nucleic Acids Res, 50, D543–D552.

105. Jumper, J., Evans, R., Pritzel, A., Green, T., Figurnov, M., Ronneberger, O., Tunyasuvunakool, K., Bates, R., Žídek, A., Potapenko, A., et al. (2021) Highly accurate protein structure prediction with AlphaFold. Nature, 596, 583–589.

