## Supplemental Figures for "Let’s make it clear: Systematic exploration of mitochondrial DNA- and RNA-protein complexes by complexome profiling"

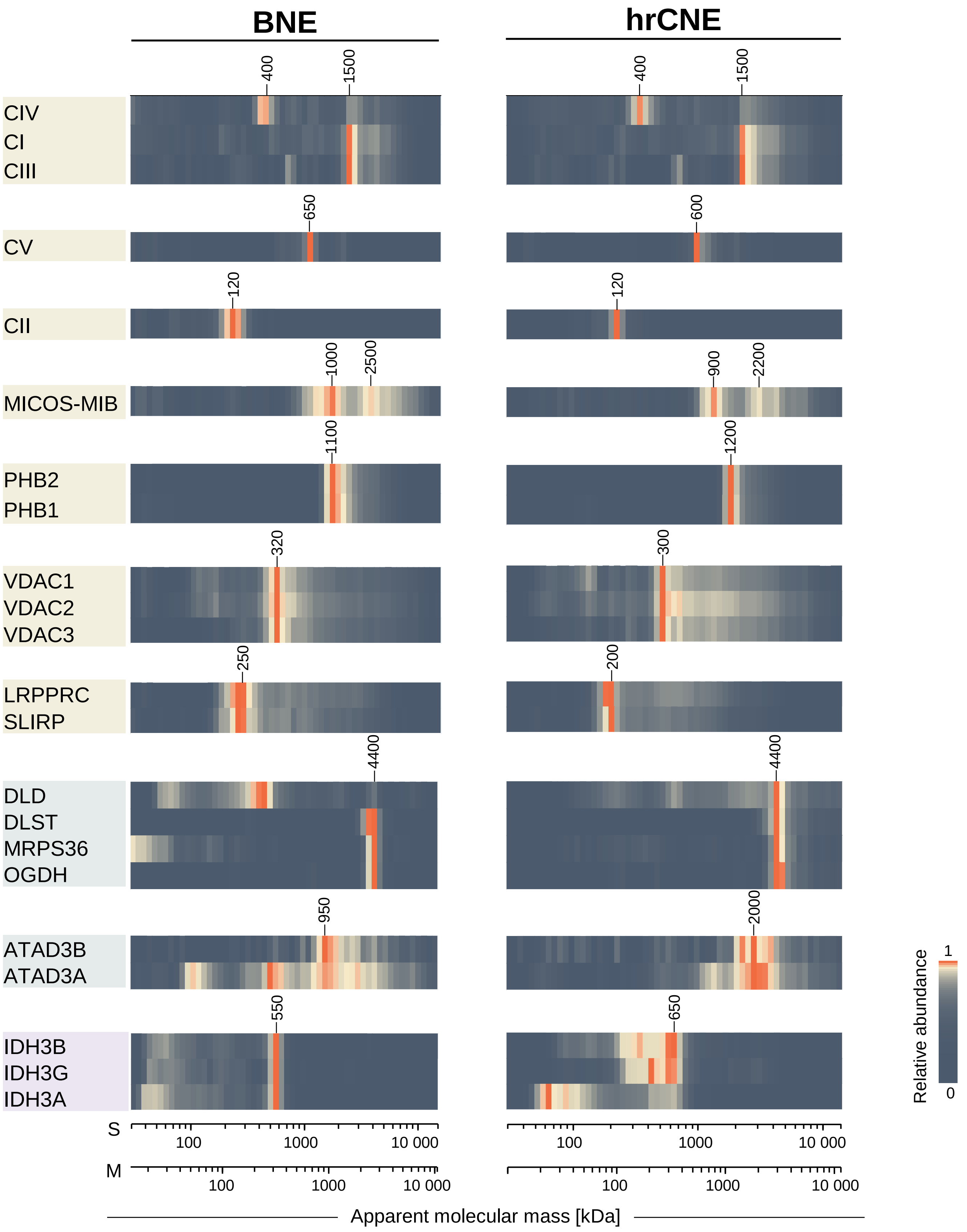


**Figure S1. Comparison of selected mitochondrial protein complexes identified in BNE- and hrCNE-based complexome profiles.** Heatmaps show relative abundances representing iBAQ values normalised to the maximum value for a given protein separately for the BNE and hrCNE profiles. Selected examples of protein complexes that were equally well detected using both protocols (yellow) or better detected using hrCNE (green). An example of a protein complex with an inconsistent migration profile using hrCNE (purple). The profiles of OXPHOS complexes (CIV, CI, CIII, CV, CII) and MICOS-MIB complex correspond to averaged profiles of all their individual subunits. Apparent molecular mass scales are provided for both soluble (S) and membrane (M) proteins.

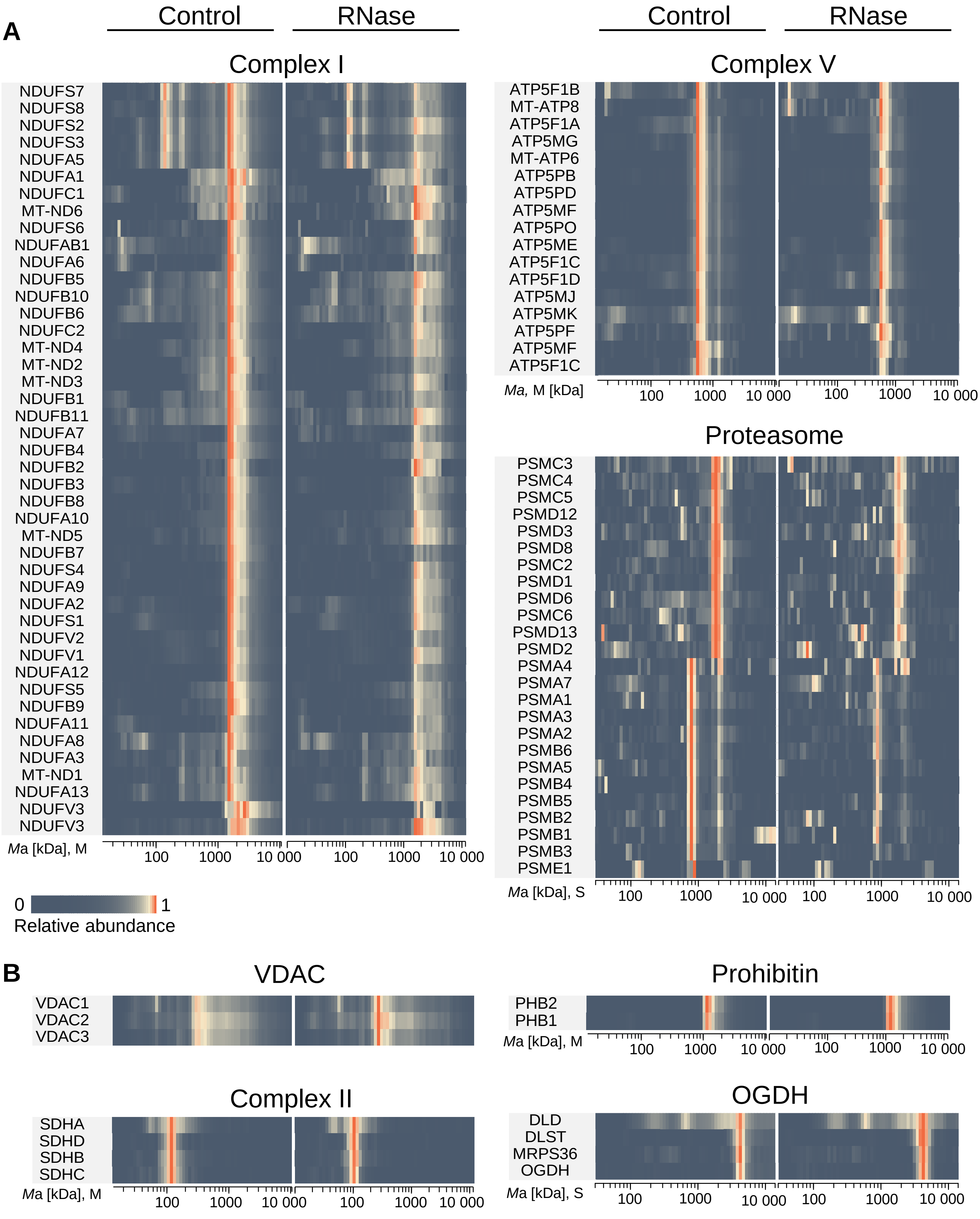


**Figure S2. Abundance profiles of selected proteins in control and RNase complexome profiles.** Heatmaps show relative abundances representing iBAQ values normalised to the maximum value for a given protein in both conditions. **A.** Components of CI, CV and proteasome displayed dissimilar patterns and/or lower intensity after RNase treatment. **B.** Reference protein complexes not affected by the RNase treatment. Apparent molecular mass scales are provided for both soluble (S) and membrane (M) proteins.

| Protein name | Gene name |
|---|---|
| 39S ribosomal protein L13, mitochondrial | **MRPL13 (LSU)** |
| Acyl-coenzyme A thioesterase 13 | ACOT13 |
| Mitochondrial import receptor subunit TOM34 | TOMM34 |
| Isoform 2 of LETM1 domain-containing protein 1 | LETMD1 |
| Aspartate aminotransferase | GOT2 |
| Peptidyl-prolyl cis-trans isomerase F | PPIF |
| Malate dehydrogenase | MDH2 |
| Probable cysteine-tRNA ligase | CARS2 |
| 28S ribosomal protein S16, mitochondrial | **MRPS16 (SSU)** |
| Malonyl-CoA-acyl carrier protein transacylase | MCAT |
| NAD-dependent protein lipoamidase sirtuin-4 | SIRT4 |

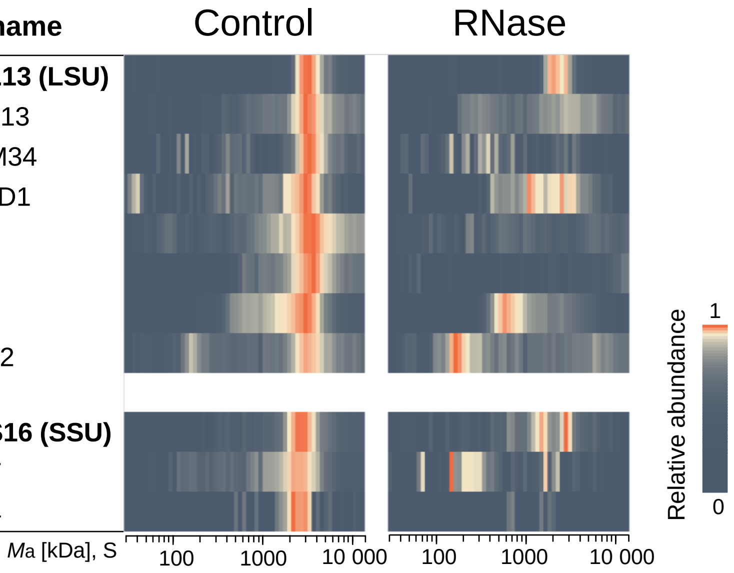


**Figure S3. Mitochondrial proteins with similar migration profiles to mitoribosomal subunits.** Segments of the control and RNase complexome profiles are shown as heatmaps. The migration of large (LSU) and small (SSU) mitoribosomal subunits are depicted by the two selected representative MRPs L13 and S16, respectively. The relative abundances were normalised to the maximum value within each pair of full profiles. The profiles represent an average of three control or two RNase independent profiles.

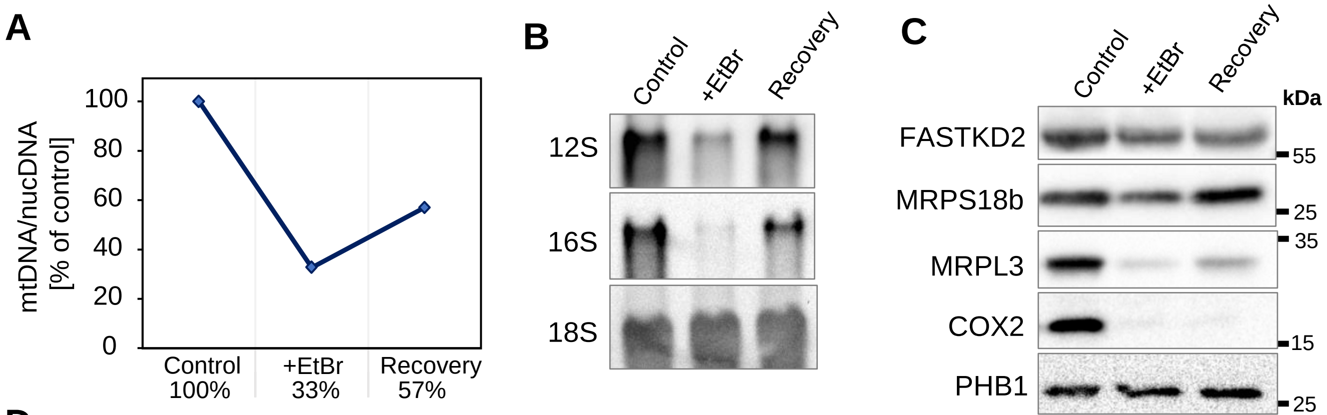


D

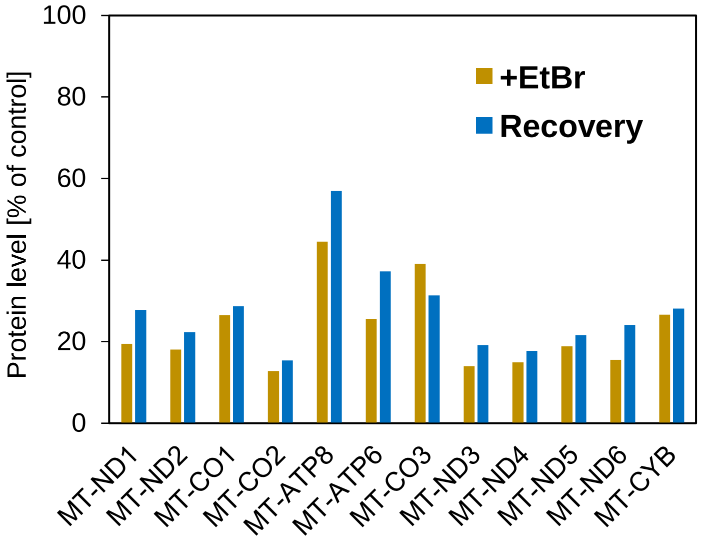


**Protein level [% of control]**

| GN | Control | +EtBr | Recovery |
|---|---|---|---|
| MT-ND1 | 100.0 | 19.5 | 27.8 |
| MT-ND2 | 100.0 | 18.0 | 22.3 |
| MT-CO1 | 100.0 | 26.5 | 28.7 |
| MT-CO2 | 100.0 | 12.8 | 15.3 |
| MT-ATP8 | 100.0 | 44.5 | 56.9 |
| MT-ATP6 | 100.0 | 25.6 | 37.2 |
| MT-CO3 | 100.0 | 39.1 | 31.3 |
| MT-ND3 | 100.0 | 14.0 | 19.2 |
| MT-ND4 | 100.0 | 14.9 | 17.8 |
| MT-ND5 | 100.0 | 18.8 | 21.6 |
| MT-ND6 | 100.0 | 15.5 | 24.1 |
| MT-CYB | 100.0 | 26.6 | 28.1 |
| Average | 100.0 | 23.0 | 27.5 |

**Figure S4. Effects of ethidium bromide (EtBr) treatment and its subsequent removal on mtDNA, mt-rRNA and mitochondrial proteins measured in the samples used for complexome profiling.** HEK293 cells were treated with 30 ng/ml of EtBr for 72 h ("+EtBr" sample). Then, EtBr was washed away and cells were grown for 24 h in EtBr-free medium ("Recovery" sample). **A.** Levels of mtDNA measured by qPCR relative to that of nucDNA. Data are shown as percentage values; the control value was set as 100%. **B**. Northern blot analysis of mitoribosomal 12S and 16S rRNA. Cytosolic 18S rRNA was used as a loading control. **C**. Levels of representative mitochondrial proteins detected by western blotting. The levels of mitoribosomal proteins S18b and L3 were reduced in response to EtBr treatment and recuperated after EtBr removal. The level of mtDNA-encoded protein COX2 was very low in both +EtBr and Recovery samples. The levels of FASTKD2 and PHB1 were not dramatically affected. **D**. The levels of mtDNA-encoded OXPHOS subunits detected by complexome profiling were estimated by summing the iBAQ values of each protein per profile. The protein levels in the +EtBr and recovery profiles are shown in both a bar chart and a table. The percentage values are relative to the control level.
